# Can the site-frequency spectrum distinguish exponential population growth from multiple-merger coalescents?

**DOI:** 10.1101/007690

**Authors:** Bjarki Eldon, Matthias Birkner, Jochen Blath, Fabian Freund

## Abstract

The ability of the site-frequency spectrum (SFS) to reflect the particularities of gene genealogies exhibiting multiple mergers of ancestral lines as opposed to those obtained in the presence of population growth is our focus. An excess of singletons is a well-known characteristic of both population growth and multiple mergers. Other aspects of the SFS, in particular the weight of the right tail, are, however, affected in specific ways by the two model classes. Using an approximate likelihood method and minimum-distance statistics, our estimates of statistical power indicate that exponential and algebraic growth can indeed be distinguished from multiple merger coalescents, even for moderate sample size, if the number of segregating sites is high enough. A normalized version of the SFS is also used as a summary statistic in an approximate Bayesian computation (ABC) approach. The results give further positive evidence as to the general eligibility of the SFS to distinguish between the different histories.

## Introduction

The site-frequency spectrum (SFS) at a given locus is one of the most important and popular statistics based on genetic data sampled from a natural population. In combination with the postulation of the assumptions of the infinitely-many sites mutation model (Watterson, 1975) and a suitable underlying coalescent framework, the SFS allows one to draw inference about evolutionary parameters, such as coalescent parameters associated with multiple-merger coalescents or population growth models.

The Kingman coalescent, developed by Kingman (1982a,b,c), Hudson (1983a,b) and Tajima (1983), describing the random ancestral relations among DNA sequences drawn from natural populations, is a prominent and widely-used model from which one can make predictions about genetic diversity. Many quantities of interest, such as the expected values and covariances of the SFS associated with the Kingman coalescent, are easily computed thanks to results by Fu (1995). The robustness of the Kingman coalescent is quite remarkable; indeed a large number of genealogy models can be shown to have the Kingman coalescent, or a variant thereof, as their limit process, cf. e.g. Möhle (1998). A large volume of work is thus devoted to inference methods based on the Kingman coalescent; see e.g. Donnelly and Tavaré (1995), Hudson (1990), Nordborg (2001), Hein *et al.* (2005) or Wakeley (2007) for reviews.

However, many evolutionary histories can lead to significant deviations from the Kingman coalescent model. Such deviations can be detected using a variety of statistical tools, such as Tajima’s *D* (Tajima, 1989a), Fu and Li’s *D* (Fu and Li, 1993) or Fay and Wu’s *H* (Fay and Wu, 2000), which are all functions of the SFS. However, they do not always allow to identify the actual evolutionary mechanisms leading to such deviations. Developing statistical tools that allow to distinguish between different evolutionary histories is, therefore, of fundamental importance.

The present work focuses on properties of the (folded and unfolded) SFS in the infinitely-many sites model for three population histories: (1) classical Kingman coalescent, (2) population growth, in particular exponential population growth, and (3) high fecundity coupled with skewed offspring distributions (HFSOD), resulting in gene genealogies being described by so-called Lambda-coalescents (Sagitov, 1999; Pitman, 1999; Donnelly and Kurtz, 1999). Briefly, multiple merger coalescents may be more appropriate for organisms exhibiting HFSOD than the Kingman coalescent (cf. eg. Beckenbach, 1994; Árnason, 2004; Eldon and Wakeley, 2006; Sargsyan and Wakeley, 2008; Hedgecock and Pudovkin, 2011), see also a recent review by Tellier and Lemaire (2014).

Recent population growth as well as multiple-merger coalescents may lead to an excess of singletons in the SFS compared to the classical Kingman coalescent based SFS, which e.g. contributes to shifting Tajima (1989b)’ *D* values to the negative. Indeed, Durrett and Schweinsberg (2005) prove that Tajima’s *D* will be negative, at least for large sample size, under fairly general multiple-merger coalescents.

The associated genealogical trees are, however, qualitatively different. While moderate fluctuations in population size lead to a time-change of the Kingman coalescent (Kaj and Krone, 2003), multiple merger coalescents by definition change the topology of the genealogical tree. There is thus hope that each demographic effect leaves specific signatures in the resulting SFS, not only with respect to an excess of singletons, but for example also with respect to its right tail.

Indeed, one observes that the Kingman coalescent will not be a good match to genetic data containing a large fraction of singleton polymorphisms (relative to the total number of polymorphisms) due to a lack of free (coalescent) parameters, as opposed to multiple merger and population growth models, which both can predict an excess of singletons. Encouragingly, multiple merger and growth models exhibit noticeable differences in the bulk of the site-frequency spectrum, in particular in the lumped tail (see Figure 1; see also (Figs. 4, S2 Neher and Hallatschek, 2013)). In Figure 1, the normalised expected spectrum 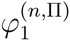 (see Eq. (2)) for a given coalescent Π, ie. the expected spectrum scaled by the expected total number of segregating sites, is compared for different multiple-merger coalescents (`B` – Beta coalescents; Schweinsberg, 2003), or (`D` – Dirac coalescents; Eldon and Wakeley, 2006), and exponential (`E`) and algebraic (`A`) growth models leading to time-changed Kingman coalescents, for sample size (number of leaves) *n* as shown. Details for these coalescent models are recalled at the beginning of the Supporting Information. The first five classes (representing relative length of external branches, two-leaf branches, etc.) are shown, with classes from six onwards collected together (labelled as ‘`5+`’). In Figure 1, the relative external branch lengths were matched between the different coalescent processes. Even though the relative external branch lengths, and by implication the number of singletons relative to the total number of segregating sites, can be matched between the different processes, the collapsed tail (group ‘`5+`’ in Figure 1) differs noticeably between the multiple-merger coalescents and the growth models. One also observes that the parameters have been chosen to match 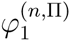 for Π ∈ {`A`, `D`, `E`} with 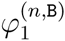 when *α* = 1, where *α* is the coalescent parameter associated with `B`. Thus 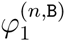 is maximized for the given *n* (since *α* ∈ [1, 2]), but 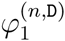, 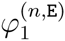, and 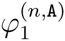 can all increase by increasing the relevant parameters (*ψ*, *β*, or *γ*).

**Figure 1:**
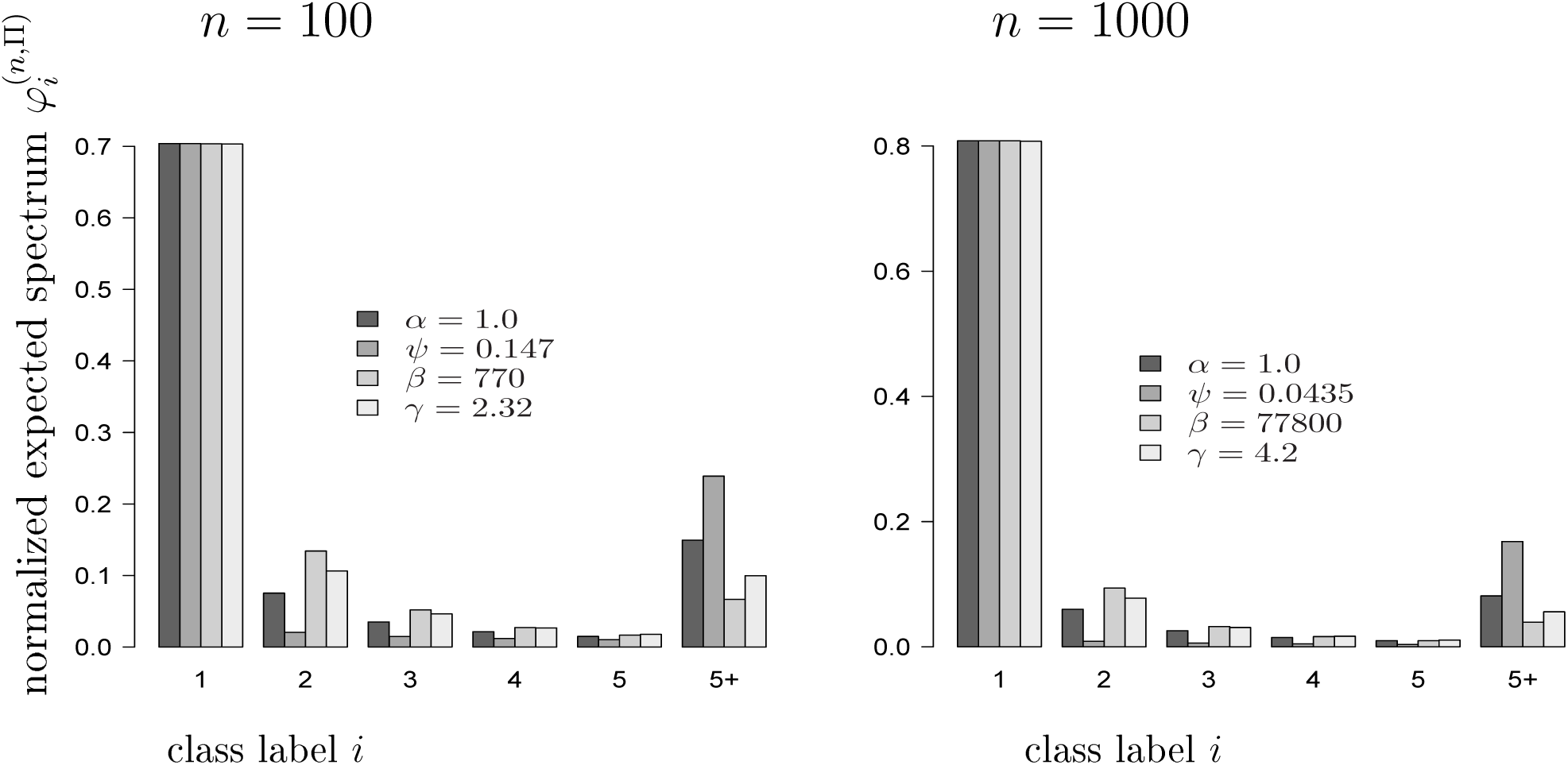
Matching 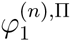 (see Eq. (2)) for the different coalescent processes Π ∈ {`A`, `B`, `D`, `E`}, with number of leaves *n* as shown. Expected values were computed exactly. The processes and their associated parameters are algebraic growth (`A`, *γ*); Beta(`2` − *α*, *α*)-coalescent (`B`, *α*); Dirac coalescent (`D`, *ψ*); exponential growth (`E`, *β*). The values with label ‘`5+`’ represent the collapsed tail 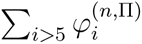.

Matching the relative external branch lengths 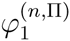 (see Eq. (2)) and observing how the rest of the normalised expected spectrum behaves, as illustrated in Figure 1, gives hope that multiple merger processes may be distinguished from (at least) particular population growth models with adequate statistical power. In the limit of large *n*, for the Kingman coalescent, 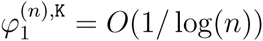.

Inference methods for distinguishing population growth from the usual Kingman coalescent have been extensively studied, see e.g. Tajima (1989a), Slatkin and Hudson (1991), Rogers and Harpending (1992), Kaj and Krone (2003) and Sano and Tachida (2005). Simulation-based work include Ramírez-Soriano *et al.* (2008), who consider the statistical power of several tests under population size increase and decrease, and the impact of recombination. Ramos-Onsins and Rozas (2002) consider the statistical power of statistics based on the site-frequency spectrum to distinguish deterministic population growth from the Kingman coalescent. On the theoretical side Myers *et al.* (2008), Bhaskar and Song (2014) and Kim *et al.* (2014) consider principal questions of identifiability of demographic histories. In particular, Bhaskar and Song (2014) show theoretically that complete knowledge of the SFS for large sample sizes carries enough information to fully recover demographic history under mild assumptions on the possible fluctations of the demography.

Detecting multiple merger coalescents in populations deviating from the Kingman coaleseent assumptions is a relatively new direction of research. Indeed, deriving inference methods based on multiple merger coalescents has only just begun (Eldon and Wakeley, 2006; Birkner and Blath, 2008; Eldon, 2011; Birkner *et al.*, 2011, 2013a,b; Steinrücken *et al.*, 2013; Koskela *et al.*, 2013; Rödelsperger *et al.*, 2014). In particular, Birkner *et al.*, (2013b) obtain recursions for the expected site-frequency spectrum associated with Lambda-coalescents. In the present work we address the issue of distinguishing multiple merger coalescents from exponential population growth, by proposing statistical tests based on the (normalized) SFS, estimating statistical power for interval hypotheses via simulation. Since we can only work with approximate likelihood functions, and our methods, in particular the so-called ‘fixed-*S*-method’, can be sensitive to an (unknown) true coalescent mutation rate *θ*/2, we complement our analysis by an approximate Bayesian computation approach (ABC; Rubin, 1984; Tavaré *et al.*, 1997; Pritchard *et al.*, 1999; Cucala and Marin, 2013; Baragatti and Pudlo, 2014).

## Theory and Methods

### Basic properties of the site-frequency spectrum

Consider a sample of *n* DNA sequences taken at a given genetic locus and assume that we can distinguish between derived (new mutations) and ancestral states. For 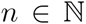 let [*n*] ≔ {1,…, *n*}. We denote by 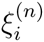 the total number of sites at which the mutant base appears *i* ∈ [*n* − 1] times. Then,

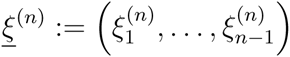

is referred to as the *unfolded* site-frequency spectrum based on the *n* DNA sequences. If mutant and wild-type cannot be distinguished, one often considers the *folded* spectrum 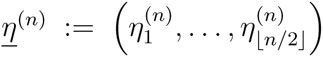, where ancestral and derived states are not distinguished, and hence

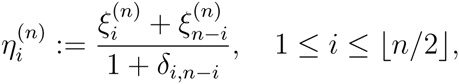

(Fu, 1995), where *δ_i_*_,*j*_ = 1 if *i* = *j*, and is zero otherwise. In this study, we will mostly be concerned with the unfolded site-frequency spectrum. Define 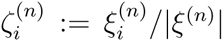 where 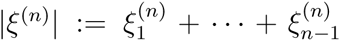 denotes the total number of segregating sites. Thus, 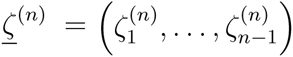 is the ‘normalized’ unfolded SFS, with the convention that 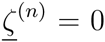 in the trivial case of complete absence of segregating sites (|*ξ*^(*n*)^| = 0).

In order to compute expected values, variances and covariances of the SFS, an explicit underlying probabilistic model is needed. In the following we assume that the genealogy of a sample can be described by a coalescent process, more precisely by either (a timechange of) the Kingman coalescent or a multiple-merger coalescent. In addition, the infinitely-many-sites mutation model (Watterson, 1975) is assumed, and mutations are modeled by a Poisson-process on the coalescent branches with rate *θ*/2. With this parametrization, the expected number of segregating sites in a sample of size 2, and hence the expected number of pairwise differences in a sample from the population, equals *θ*.

Closed-form expressions for the expected values and (co)variances of 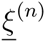 have been determined in Fu (1995) when associated with the Kingman coalescent. One can represent the expected values of 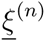 in a unified way using the results of Griffiths and Tavaré (1998), Kaj and Krone (2003) and Birkner *et al.* (2013b), that allow to treat the expected values (and covariances) of the SFS for all coalescent models in question.

Let 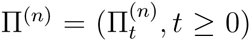 be a (partition-valued exchangeable) coalescent process started from *n* leaves (partition blocks) corresponding to the random genealogy of a sample of size *n*. By discussing ‘leaves’ rather than DNA sequences we are emphasizing our viewpoint of the genealogy as a random graph, where the leaves are a particular kind of a vertex. Our interest is in the topology of the genealogy and how it is reflected in the associated site-frequency spectrum.

If the initial number of leaves is not specified, we simply speak of Π. One may think of Π as the Kingman coalescent, but the point is that the following result will stay true also for externally time-changed Kingman coalescents as well as multiple merger coalescents (a.k.a. ‘Lambda’- or ‘Xi’-coalescents in the mathematical literature), and even externally time-changed multiple merger coalescents.

Given *n* and a coalescent model Π, let 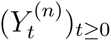 be the block counting process of the underlying coalescent Π^(*n*)^ started from *n* lineages, i.e. 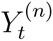 gives the number of ancestral lines (blocks) present/active at (backwards) time *t*. For 2 ≤ *k* < *n*, let 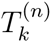 be the random amount of time that 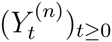 spends in state *k*. Given a coalescent Π^(*n*)^ started from *n* (unlabelled) lineages, denote by *p*^(*n*),Π^[*k*, *i*] the probability that, *conditional* on the event that 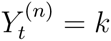 for some time point *t*, a given one of the *k* blocks subtends exactly *i* ∈ [*n* − 1] leaves. A general representation of 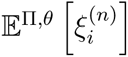 is then

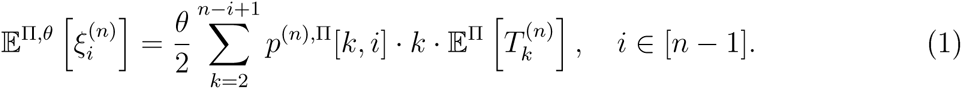

The normalized expected SFS, 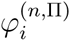, for *i* ∈ [*n* − 1], is defined as

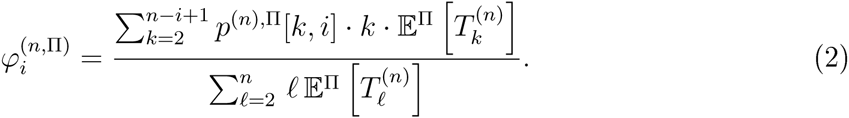

where the denominator in (2) is the expected total tree length when starting from n leaves. One can interpret the quantity 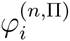 as the probability that a mutation, under the infinitely many sites assumption and the coalescent model Π, with known ancestral types, appears *i* times in a sample of size *n*. Importantly, 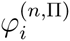 is not a function of the mutation rate, unlike 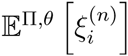. One can also view 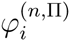 as a first-order approximation of the expected value 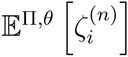 of the normalised SFS.

As examples for Π we will consider the classical Kingman coalescent (`K`), exponential (`E`) and algebraic (`A`) growth models, and the Beta(2 − *α*, *α*) (`B`) and the Dirac (`D`) multiple-merger coalescents, as recalled in the Supporting Information. Simulations suggest that 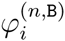 is a good approximation of 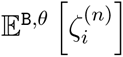 when *α* is not too close to 1, and *n* and *θ* are not too small (Birkner *et al.*, 2013b). Similar conclusions hold in the case of exponential and algebraic growth (results not shown).

One can use the recursive formulas obtained by Birkner *et al.* (2013b) to compute 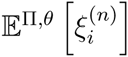 associated with Lambda-coalescents. To compute 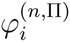 associated with growth models we use the results of Polanski and Kimmel (2003), whose recursions are recalled in the Supporting Information.

A comparison of the observed 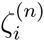 (instead of 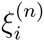) to an expected value 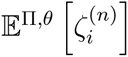 – obtained under a particular coalescent model Π – enables one to do inference without having to jointly estimate the mutation rate *θ* using e.g. a minimum-distance statistic. Indeed, it appears that under any coalescent model Π, 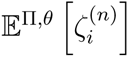 is almost constant as a function of the mutation rate *θ* (unless *θ* is very small); we provide some evidence for this in the Supporting Information, see (S12). Unfortunately, there seems to be no explicit way of representing 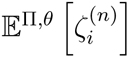 as a simple function of the associated coalescent parameters and sample size *n*. As mentioned above, one may instead work with 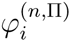.

### Timescales, segregating sites and mutation rates

The choice of a multiple merger coalescent model (resp. demographic history) Π and its underlying parameters strongly affects classical estimates for the coalescent mutation rate *θ*/2 (i.e. the Poisson rate at which mutations appear on coalescent branches). Assume w.l.o.g. for all multiple merger coalescents in question that the underlying coalescent measure Λ is always a probability measure: This normalisation fixes the coalescent time unit as the expected time to the most recent common ancestor of two individuals sampled uniformly from the population.

Given an observed number of segregating sites *S* in a sample of size *n*, a common estimate 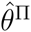 of the scaled mutation rate *θ* associated with coalescent model Π is the Watterson estimate

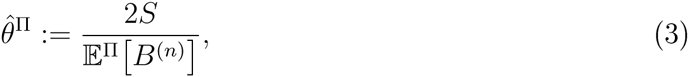

where 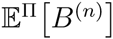 is the expectation of the total tree length *B*^(*n*)^ of an (*n*-) coalescent model Π. One can of course also estimate *θ* as a (different) linear combination of the site-frequency spectrum [(cf. Achaz, 2009) in the case of the Kingman coalescent]. Using the recursions for 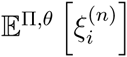 obtained by Birkner *et al.* (2013b), one can also estimate *θ*, using either (3) or a linear combination of the expected SFS, in case of a Lambda-coalescent.

Given an estimate 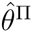, and knowledge of the mutation/substitution rate 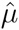 per year at the locus under consideration, allows one to find a real-time embedding, of the coalescent history via the approximate identity

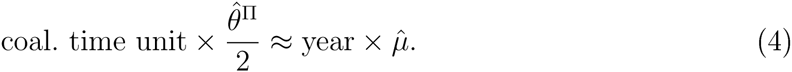

cf. (Steinrücken *et al.*, 2013, Section 4.2), which of course depends on Π.

If one has additional information on the specific reproductive mechanisms of an approximating population model, this can even enable one to estimate the model census population size. For example, given a Cannings population model (Cannings, 1974, 1975) of fixed size *N*, let *c_N_* be the probability that two gene copies, drawn uniformly at random and without replacement from a population of size *N*, derive from a common parental gene copy in the previous generation. While for the usual haploid Wright-Fisher model *c_N_* = 1/*N*, in a class of population models studied by Schweinsberg (2003) leading to the Beta(2 − *α*, *α*)-coalescent, *c_N_* is proportional to 1/*N^α^* ^− 1^, for *α* ∈ (1, 2] (but note that the proportionality constant depends on finer details of the particular model). By a limit theorem for Cannings models of Möhle and Sagitov (2001), one coalescent time unit corresponds to approximately 1/*c_N_* generations in the original model with population size *N*, Thus the mutation rate 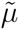 at the locus under consideration per individual per generation must be scaled with 1/*c_N_* (as noted e.g. in Eldon and Wakeley (2006)), and the relation between 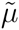, the coalescent mutation rate *θ*^Π^/2 and *c_N_* is then given by the (approximate) identity

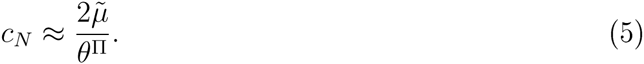

In particular, if the Cannings model class (and thus *c_N_* as a function of *N*) is known, *N* can be estimated via (5).

In this context, it is important to note that different population models on very different timescales can still have the Kingman coalescent as their ancestral limit process; two examples are the Wright-Fisher 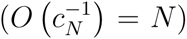 and the Moran 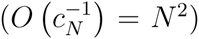 models. This is certainly also the case for multiple merger coalescents. In particular, *c_N_* is a priori *not* a function of the limiting coalescent model (this appears to be a rather frequent misperception).

Distinguishing real- and coalescent-timescales is important, because non-linear scaling may otherwise easily lead to confusion: For example, the expected total tree length 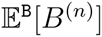 (measured in coalescent time units) *decreases* as a function of *α* ∈ (1, 2], while the corresponding quantity (measured in real-time generations) 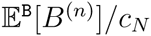 *increases* in *α* ∈(1, 2] (cf. Figure S1 in the supporting information).

The time-scaling applied to a classical Wright-Fisher model with *fluctuating* population size [as in Kaj and Krone (2003)] in order to obtain a (time-changed) Kingman coalescent is recalled in the Supporting Information, see in particular Equation (S2), Again, the estimate (3) of *θ* depends on the growth model and growth parameter.

### Approximate likelihood ratio tests for the SFS

Our aim is to construct a statistical test to distinguish between the model classes `E`, `A`, `D` and `B` (which intersect exactly in the Kingman coalescent `K`). In order to distinguish, say, `E` from `B`, based on an observed site-frequency spectrum 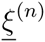 with sample size n and segregating sites 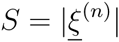, a natural approach is to construct a likelihood-ratio test.

Recall that we think of our observed spectrum as a realization of a coalescent tree with *n* leaves obtained from a coalescent model Π, with mutations distributed on the tree according to an independent Poisson(*θ*/2)-process. For each model Π from classes {`E`, `A`, `D`, `B`}, the coalescent will be uniquely determined by a single coalescent parameter *β* ∈ [0, ∞) (for 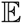), *γ* ∈ [0, ∞) (for `A`), *ψ* ∈ [0, 1] (for `D`) and *α* ∈ [1, 2]. (for `B`)^1^.

Suppose our null-hypothesis *H*_0_ is presence of recent exponential population growth (`E`) with (unknown) parameter *β* ∈ [0, ∞), and we wish to test it against the alternative *H*_1_ hypothesis of a multiple merger coalescent, say, the Beta(2 − *α*, *α*)-coalescent (`B`) for (unknown) *α* ∈ [1, 2], where *β* = 0 and *α* = 2 correspond to the Kingman coalescent. In this framework, the coalescent mutation rate *θ* is not directly observable, but plays the role of a nuisance parameter. In particular, it is the interplay of the coalescent model Π and the mutation rate parameter *θ* that governs the law of the observed number of mutations, cf. the discussion in the previous section. To take *θ* explicitly into account, one could test

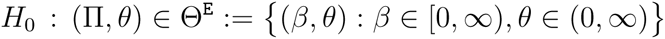

(exponential growth) against

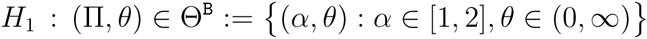

if the Beta-coalescent family is the alternative (by slight abuse of notation, we identify the coalescent model Π with the corresponding coalescent parameter *β* (resp. *α*) in each model class when appropriate). The underlying parameter ranges are two-dimensional; and although an explicit likelihood-ratio test based on methods described in Simonsen *et al.* (1995) can be constructed, it will likely pose computational challenges.

Instead, given an observed number of segregating sites *S* = *s*, we simplify our framework by employing the ‘fixed-*s*-method’ discussed e.g. in Depaulis and Veuille (1998) and Ramos-Onsins and Rozas (2002). Here we treat the observed number of segregating sites as a *fixed parameter* 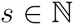, not as (observation of a) random variable *S*. We will thus obtain the empirical distributional quantities of our test by Monte-Carlo simulations, placing uniformly at random *s* mutations along the branches of the simulated tree.

The fixed-*s*-method is different from generating samples for a given *θ* by conditioning on *S* = *s*, yet the fixed-*s*-method usually leads to reasonable tests when the true *θ* is close to the Watterson estimate 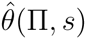 based on *s* [Wall and Hudson (2001)]. However, it can lead to substantial deviations from the condional distribution if *θ* is extreme (see e.g. Markovtsova *et al.* (2001), who show that for a test in a related framework, the probability of rejection can be substantially different from 5%). Regarding this caveat, Depaulis *et al.* (2001) take a Bayesian viewpoint and show that those values of *θ* that lead to unreliable tests are highly unlikely given *s*. We address this issue by using rejection sampling to check the robustness of the fixed-*s*-method against varying *θ* (see Supporting Information). Our analysis is complemented with an ABC approach using the normalized frequency spectrum 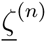, which should be insensitive to the actual value of *θ* as long as *θ* is not too small, cf. (S12).

By fixing *S* = *s* and treating it as a parameter of our test, we may consider the new pair of hypotheses

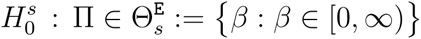

and

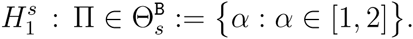

Define a likelihood-function *L*(Π, <u>*k*</u>^(*n*)^, *s*) for the observed frequency spectrum 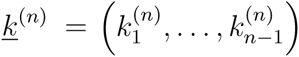 with fixed |<u>*k*</u>^(*n*)^| = *s* under the coalescent model 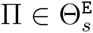 (resp. 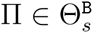), by

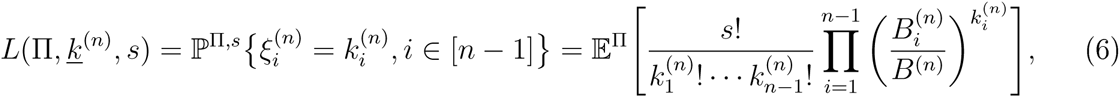

where the 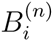 are the random length of branches subtending *i* ∈ [*n* − 1] leaves and *B*^(*n*)^ is the total branch length of the coalescent under Π. The fixed-*s*-paradigm thus leads to a mixture of multinomial distributions, where the parameters are given by the respective relative branch lengths. The hope is that the location of the maximum of *L*(Π, <u>*k*</u>^(*n*)^, *s*) is typically not far from the location of the corresponding coordinate of the maximizer in the full two-dimensional ‘explicit-*θ*’-model, in which one can additionally maximize over all *θ* ∈ [0, ∞).

Now we can construct a ‘likelihood-ratio’ test based on *L*(Π, <u>*k*</u>^(*n*)^, *s*) via the likelihood-ratio function

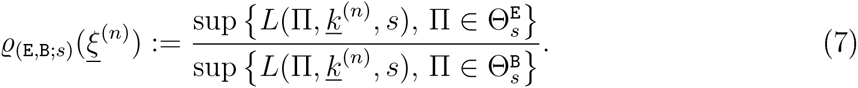

Given a significance level *a* ∈ (0, 1) (say, *a* = 0.05), let 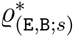 (*a*) be the *a*-quantile of 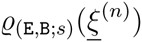 under `E`, chosen as the largest values so that

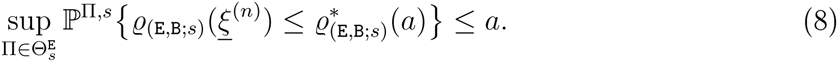

The decision rule that constitutes the ‘fixed-*s*-likelihood-ratio test’, given *s* and sample size *n*, is

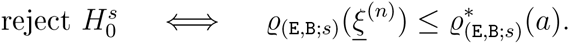

This formulation is free of the nuisance parameter *θ*, To assess the justification of the ‘fixed-*s*-assumption’, we investigate how close this is to the corresponding quantiles for different values of *θ*, including the Watterson estimator

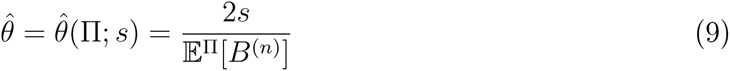

cf. (3), for selected choices of Π. The agreement appears reasonably good and seems to increase with sample size, cf. the supplementary material.

The corresponding power function of the test, that is, the probability to reject a false null-hypothesis, is given by

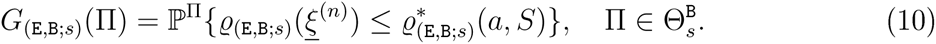

The likelihood (6) cannot be represented as a simple formula involving the coalescent parameters; one can approximate (6) via a Monte Carlo approach but this is computationally expensive. An approximation is

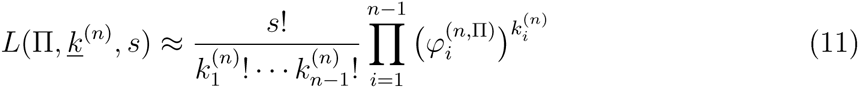

where we replaced the random quantities 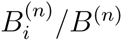 in (6) by 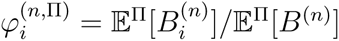 (2).

Interestingly, an approximate maximum likelihood method based on (11) is equivalent to the following approach: Consider a family of (approximate) likelihood functions

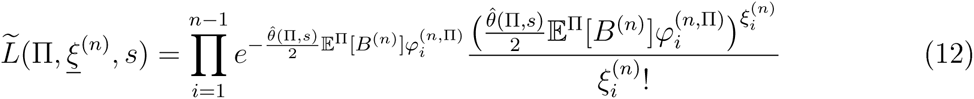

where 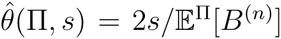 is the Watterson estimator for the mutation rate under a Π-coalescent with *n* leaves when *S* = *s* segregating sites are observed, recall (3). In (12), 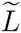 is well defined even if 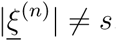.

The rationale behind (12) is simple: It pretends that the classes are approximately independent and Poisson-distributed (this is of course not literally true but encouraged by the fact that the off-diagonal entries of the covariance matrix of 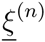 are small compared to the diagonal terms, see Birkner *et al.* (2013b); our approximation is different from Sawyer and Hartl (1992)’s Poisson random field which applies to unlinked sites).

For fixed *s*, we can view 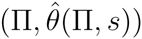 as parametrizing a one-dimensional curve in the full two-dimensional space *H*_0_ ∪ *H*_1_ defined by the requirement that 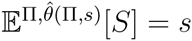.

The two approximate maximum likelihood approaches based on (11) and on (12) are equivalent; indeed

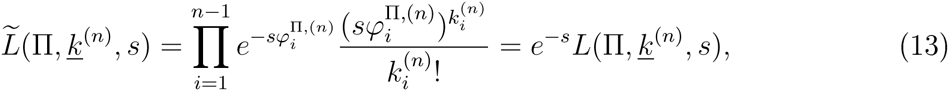

since the 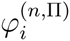 sum to 1. Hence, both likelihood functions differ only by the fixed prefaetor *e*^−*s*^, so that they attain their maximum at the same position.

Thus, now we consider the statistic

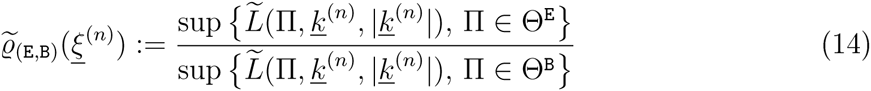

(where Θ^E^ and Θ^B^ refer to the projection of *H*_0_, respectively *H*_1_, on the coalescent parameter). For a given value of *s*, we can then (by simulations using the fixed-*s*-approach) determine approximate quantiles 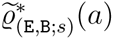 associated with a significance level *a* as in (8), and base our test on the criterion 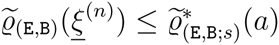. Similary, the (approximate) power function 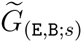 can be estimated using simulations.

An alternative approach to the (approximate) likelihood-based tests would be rejection rules based on minimal distance statistics:

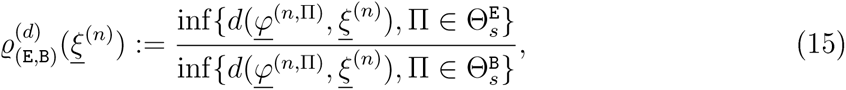

for some suitable distance measure *d* (e.g. the **ℓ*_p_* distance with *p* = 2) with corresponding power function 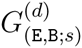. We will not discuss the theoretical justification of this method. However, we will use the **ℓ**_2_-distance between normalized expected spectra under various coalescent models to produce three-dimensional ‘heatmaps’ that give some intuitive insight on how a pair of different models out of {`E`, `B`, `A`, `D`} relates to each other depending on the underlying pair of coalescent parameters (cf. Results).

We conclude this section by a remark on *lumping*. One often observes 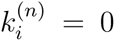 for most *i* greater than some (small) number *m* in observed data, in particular for large *n*. It thus seems natural to consider (approximate) likelihood functions for ‘lumped spectra’ (e.g. collapsing all entries in classes to the right of some number *m* into one class *m*^+^), as we have done eg. in Figure 1. Another natural type of lumping may be to collect together classes so that 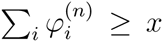 for some *x* ∈ (0, 1/2]. This may not always be feasible, though, if the individual 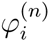 quickly become quite small, and we will refrain from going into a more detailed theoretical discussion of optimal lumpings. However, we will see in our subsequent ABC analysis, that adequate lumping can improve the reliability of our model-selection procedure.

### Approximate Bayes factors and model selection

In view of the approach and notation of the previous section, an analogous way of model selection could be based on a *Bayes factor* of the form

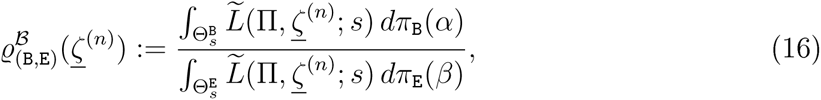

(and similar for all other combinations of classes `A`, `D`, `E`, `B`), given a pair of priors *π*_B_, *π*_E_ on 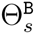, 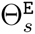. While the approach will work in principle also for the two-dimensional prior ranges Θ^B^ and Θ^E^, we will present the (approximate) Bayesian methods with one-dimensional prior ranges (where *S* = *s* is treated as a fixed parameter, motivated from the ‘fixed-*s*-approach’), so that they complement our previous methods.

Our simulations will be obtained using the rationale behind (12), that is, after simulating a tree according to a given coalescent parameter, say *α* from *π*_B_, mutations are placed on the tree according to a Poisson process with mutation rate 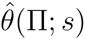 estimated using (3). However, in (16), we use the normalized site frequency spectrum 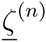 as observed statistics, since it should be more insensitive to the ‘true’ coalescent mutation rate *θ* [potentially deviating from 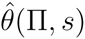], as argued in the corresponding section in the Supporting Information, and thus yield more robust results. 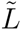 in Eq. 16 thus denotes the likelihood function of the observed nSFS under the chosen coalescent model with mutation parameter given by Watterson’s estimator based on *s*. Since we estimate the mutation rate based on *s*, the information loss of using the nSFS instead of the SFS should be only slight. We also experimented with ABC based on simulations using the fixed-*s*-method, i.e. distributing a fixed number of mutations uniformly on the simulated tree, and generally found higher misclassification probabilities (results not shown).

To overcome the problem of exact computation of 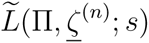, which appears infeasible in practice, we employ approximate Bayesian methods (see e.g. Beaumont (2010)) based on the *ℓ*_2_-distance between observed and simulated nSFS.

Bayes factors based on further (lumped) distances *d* and/or the folded nSFS may of course also be considered. In line with classical Bayes factor philosophy (cf. e.g. Kass and Raftery (1995)), one interprets an observed value of 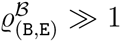 as evidence in favor of 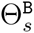 over 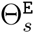.

For the ABC analyses, we consider as before on exponential growth (`E`), algebraic growth (`A`), and Beta-(`B`), and Dirac-(`D`) coalescents. Given sample size *n* and number of segregating sites *s*, again the coalescent model classes can be parametrised by a single parameter each, which are the exponential growth rate *β* ∈ [0, ∞), algebraic growth rate *γ* ∈ [0, ∞); Beta coalescent parameter *α* ∈ [1, 2], and mass point location *ψ* ∈ (0, 1] for the Dirac coalescent.

For convenience, we employ a simple rejection-based ABC scheme to approximate the Bayes factor for the model (class) comparison given an observed nSFS (resp. folded and/or lumped versions, which can be treated analogously). First, select a number of models (out of {`E`, `B`, `A`, `D`}) that should be compared, say `E` and `B`, and choose the corresponding prior distributions on the correspoding coalescent parameter ranges. To simulate say *n*_r_ independent samples of the nSFS from each model, say from `E`, independently generate *n*_r_ coalescent parameters from the prior *π*_E_, and a corresponding coalescent tree Π for each generated coalescent parameter. Distribute independently Poisson mutations with parameter 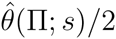 on each such tree and record the corresponding normalized site frequency spectra. This should be done independently for all models.

Then fix a tolerance level *x* ∈ (0, 1) and count the number of simulations *N*_E_, *N*_B_ from each model that are among the 100 · *x*% best fits with respect to the *ℓ*_2_ distance to the observed nSFS 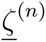 (the ‘accepted’ simulations). Here, we use an additional scaling by dividing each class (resp. lumped class) in the nSFS by the median (if non-zero) within this class observed in all simulations as implemented in the R package abc (Csilléry *et al,* 2012), The Bayes factor for model `E` vs. `B` can then be approximated by

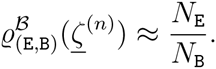

To assess how well this ABC approach allows one to distinguish, say, `E` from `B` (or, more generally, simultaneously among {`E`, `B`, `A`, `D`}), we use two approaches from the `R` package `abc`. Both are based on leave-one-out cross-validation. More precisely, we pick *n*_cv_ simulations at random from each model, treat them as the observed value of the nSFS and then run the ABC approach with the same parameters and simulations as above. For each crossvalidation sample, say 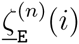, *i* ∈[*n*_cv_] from model `E`, we record the counts of accepted simulations 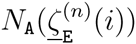, 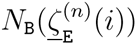 and 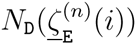 from the model classes `A`, `B`, `D`, `E` (recall that the chosen cross-validation sample is left out). As measures for the distinction ability of this approach, we record for each model class, borrowing notation from Stoehr *et al.* (2014),

- the (estimated) mean posterior probabilities *π* for model `B` given the observed nSFS under the true model `E`, say,

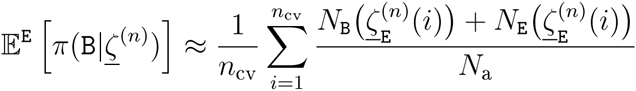

where 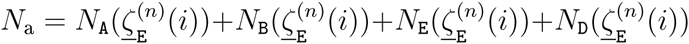 is the number of accepted simulations;
- the (estimated) mean misclassification probabilities

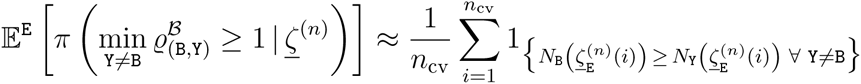

for `Y` ∈ {`A`, `E`, `D`}. To ease notation we will from now on omit *n* in the formulae.

In practice, we need to efficiently generate samples of the nSFS under the different models which can be achieved by backward-in-time coalescent simulations. For the exponential growth models (`E`), we use Hudson’s ms (Hudson, 2002) as implemented in the `R` (R Core Team, 2012) package phyclust (Chen, 2011). For algebraic growth models, the Beta-coalescents (`B`) and Dirac coalescents (`D`), we use custom `R` and `C` scripts to generate samples of the nSFS, available at http://page.math.tu-berlin.de/}eldon/programs.html. To conduct the actual ABC analysis including cross-validation techniques, we employed the `R` package abc (Csilléry *et al.* (2012)). Additionally, since we use Watterson’s estimator to set the mutation rate within each model, we compute the mean total length of each coalescent model as described in Supporting Information.

## Results

### Power estimates of approximate likelihood-ratio tests

To assess the sensitivity of our approximate likelihood ratio test associated with the likelihood ratio function (7), we estimate its power 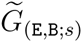 from the analogue of (10) based on the approximate likelihood from (12) as a function of *α* (Figure 2A) with 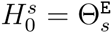 and 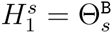; and estimate 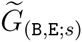 as a function of *β* with 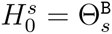 and 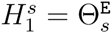 (Figure 2B).

**Figure 2:**
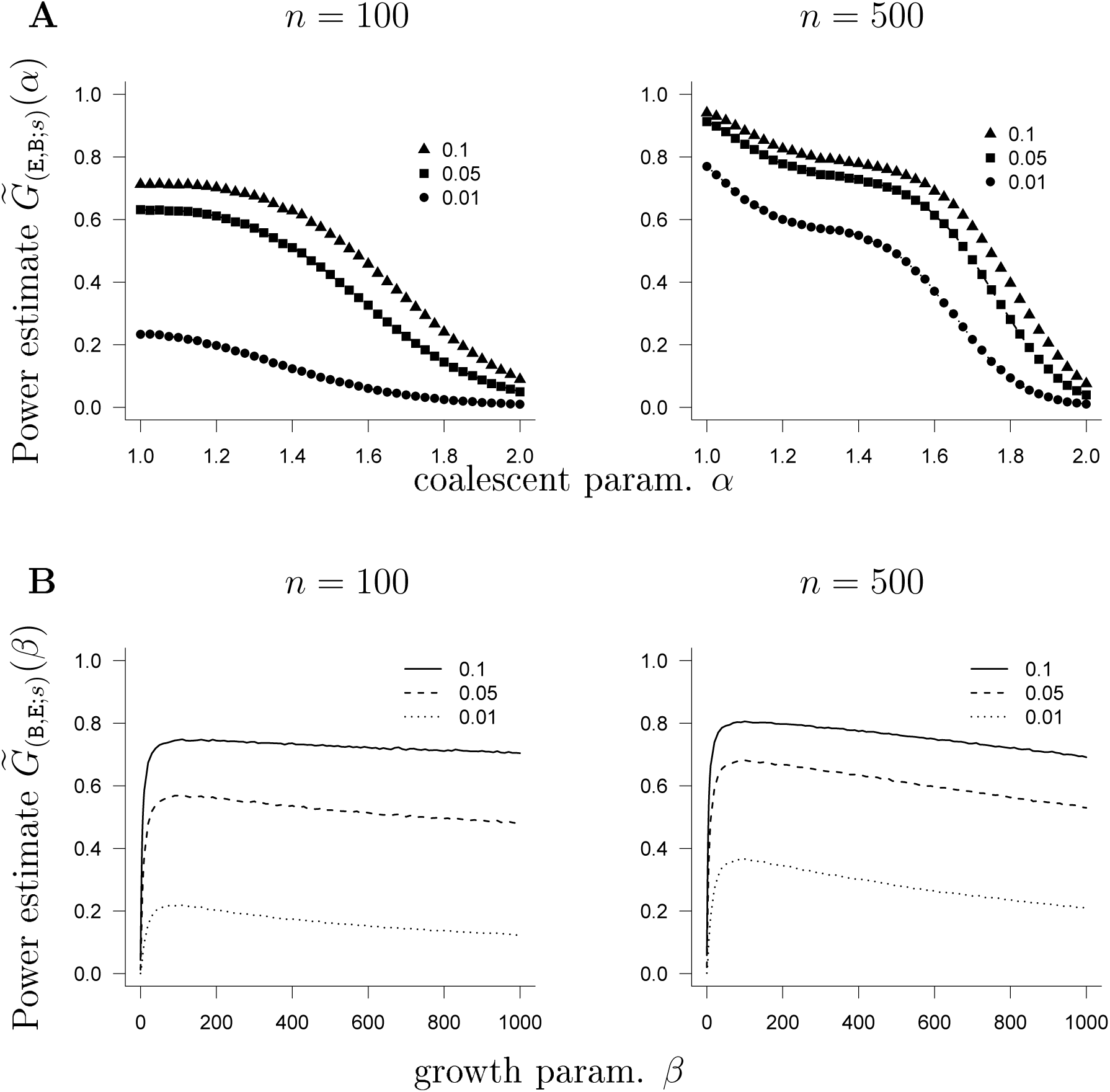
**A**: Estimate of 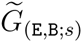 from (10) based on the approximate likelihood from (12) as a function of *α* (no lumping) with number of leaves *n* as shown, and *s* = 50. **B**: Estimate of 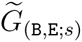 from (10) based on the approximate likelihood from (12) as a function of *β* (no lumping) with number of leaves *n* as shown, and *s* = 50. The symbols denote the size of the test as shown in the legend. The interval hypotheses are discretized to 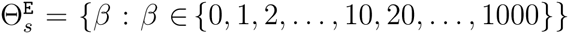 and 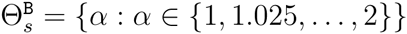. In **A**, the Beta(2 −*α*, *α*)-coalescent is the alternative; in **B** exponential growth is the alternative.

As shown in Figure 2, reasonably high power is obtained to reject 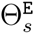 for *n* = 500; and even for smaller sample size *n* = 100, but the power also depends, as one would expect, on the size of the test. As a sidenote, we remark that the power estimates 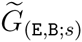, as a function of *α*, are right at the size of each corresponding test when *α* = 2 (the Kingman case), as required.

The mtDNA-genome analysis of Carr and Marshall (2008), who scanned whole mitochondrial genomes (15,655 bp) of the highly fecund Atlantic cod (*Gadus morhua*), prompted us to briefly investigate the power (Figures S3, S4) with number of segregating sites *s* = 300. This is nearly the total number of polymorphisms (298) observed among the 32 mtDNA genomes sampled by Carr and Marshall (2008). Our results show that while we may not quite have enough power when *n* = 30 and *s* = 300, (Figure S4A), we would be in good shape for *n* = 100 (Figures S3, S4B). It would be very interesting to analyse such a sample once available, since it appears to be an open debate whether Beta-coalescents should be favoured over classical models (including recent population growth) in HFSOD populations (cf. eg. Steinrücken *et al.* (2013)).

Another quite striking observation is that the power of our test is apparently nonmonotone as a function of *β* when 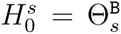, in particular for smaller Type I error. We will present a possible heuristic explanation for this in the discussion section.

Rather high power in general is obtained when comparing 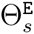 and 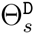 associated with the Dirac coalescent (Figure S2) for (*n*, *s*) = (200, 50).

For further combinations, we refer to the supporting material, where 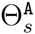, associated with algebraic growth, is compared with 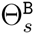 in Figure S6, and with 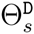 in Figure S5. The power functions 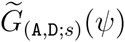 (Figure S5) are decidedly non-monotoe, as is 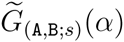 (Figure S6A).

We conclude with a short remark on the sensitivity of our results on *lumping* of classes in the observed spectrum. Indeed, our power estimates suggest that keeping at least the first five classes of the SFS intact, and collecting the rest into one other class, has little effect on the power of the test (results not shown). Keeping only the singleton 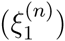 class intact, and collecting all the rest into one class, however, significantly diminishes power (results not shown).

*Availability of source-code:* `C` (cf. Kernighan and Ritchie, 1988) code written for estimating the power of our tests, where use was made of the GNU Scientific Library (Galassi *et al.*, 2013), is available at http://page.math.tu-berlin.de/}eldon/programs.html.

### Mean-squared distance landscapes for the normalized expected SFS under different growth and coalescent models

Given the potential ability to distinguish between growth- and multiple-merger coalescent models invites the following question: How does the distance between 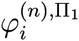 and 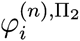 behave as a function of the underlying coalescent and growth parameters? Is it possible to visibly identify a one-dimensional curve given by coalescent parameter pairs corresponding to (Π_1_, Π_2_) along which minimal distance is achieved? Figures 3 and 4 are a brief effort to understand the relation between the normalised expected SFS for the models in question by graphing the *ℓ*_2_-distance 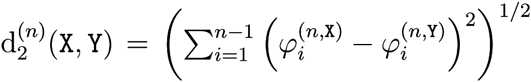 as a function of the coalescent and growth parameters associated with `X` and `Y`. In Figure 3 `E` is compared to `B` and `D`. In Figure 4, `A` is compared to `B` and `D`. In Figures 3 and 4, the upper panels show the distance as the respective growth parameter ranges from 0 to 1000, while the lower panels ‘zoom in’ on the range from 0 to 10.

**Figure 3:**
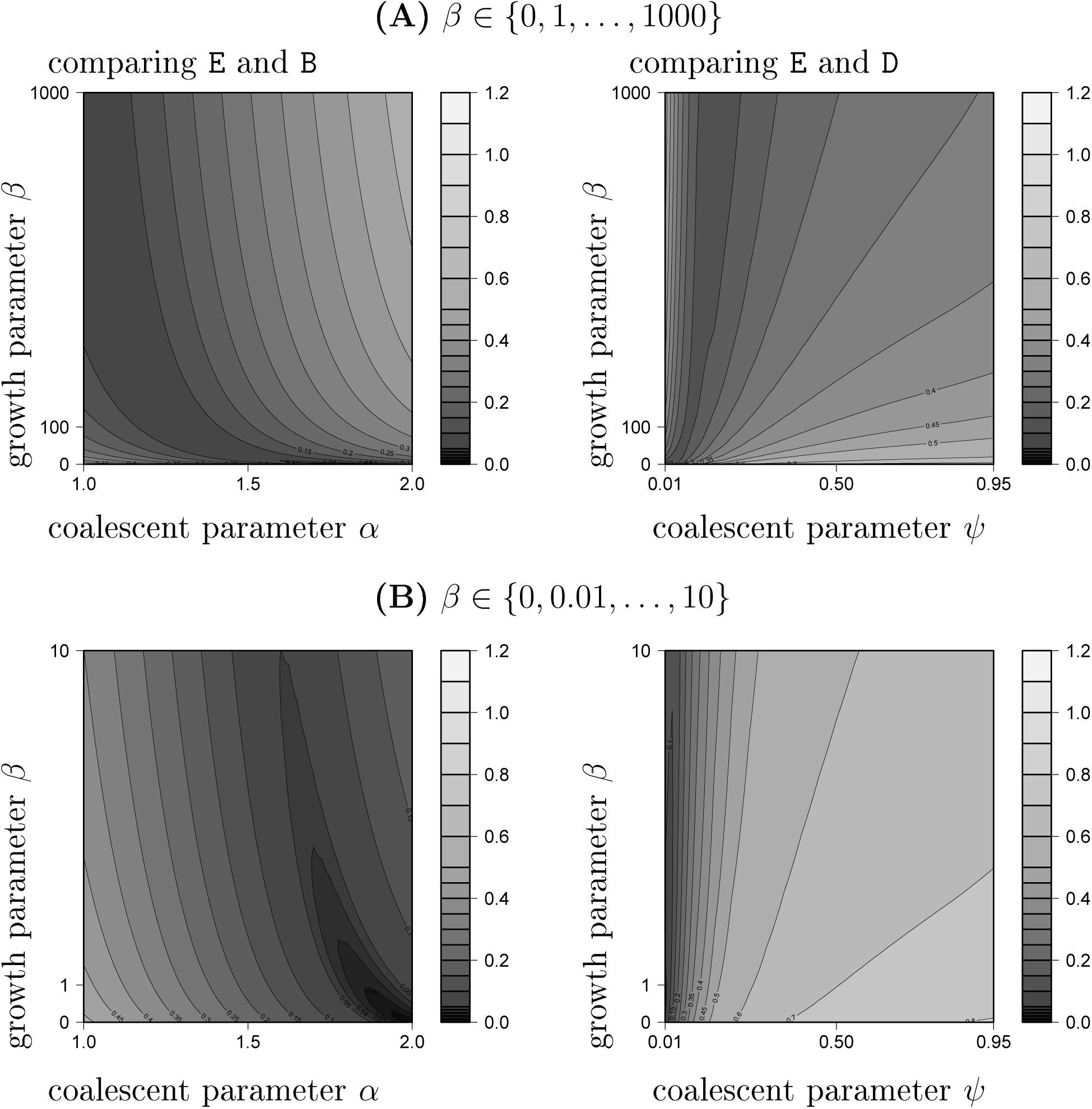
The *ℓ*_2_-distance 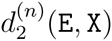 for `X` ∈ {`B`, `D`} of the normalized expected spectra 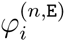 (see Eq. (2)) and 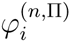 as a function of *α* (`X` = `B`) (resp. *ψ*; `X` = `D`) and *β* `E` for number of leaves *n* = 100. Expected values were computed exactly. The gridpoints are *α* ∈ {1, 1.025,…, 2}, *ψ* ∈ {0.01, 0.02,…, 0.1, 0.15, 0.2,…, 0.95}, and for *β* as shown.

**Figure 4:**
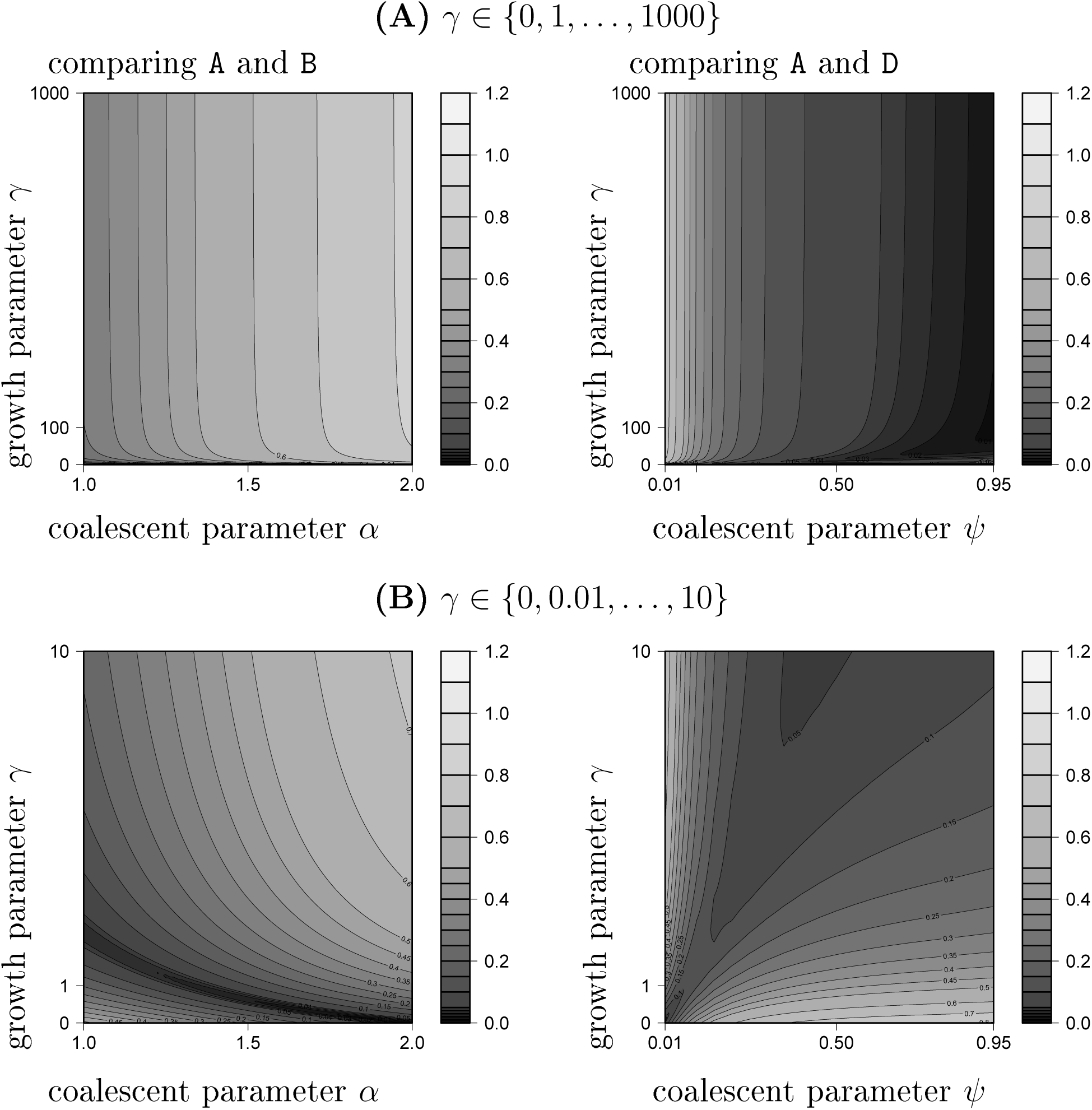
The *ℓ*_2_-distance 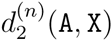 for `X` ∈ {`B`, `D`} of the normalized expected spectra 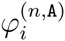 (see Eq. (2)) and 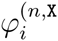 as a function of *α* (`X` = `B`) (resp. *ψ*; `X` = `D`) and *γ* (`A`) for number of leaves *n* = 100. Expected values were computed exactly. The gridpoints are *ψ* ∈ {0.01, 0.02,…, 0.1, 0.15, 0.2,…, 0.95}, *α* ∈ {1, 1.025,…, 2}; for *γ* as shown.

Figure 3 indicates the presence of a region, essentially a curve in the two-dimensional (*α*, *β*) parameter space, along which the lowest *ℓ*_2_ distance is reached. However, one should be aware that this curve shifts in space when the sample size *n* is increased (data not shown).

Figure 3 suggests that we should have good power to distinguish between algebraic growth and Beta-coalescents. However, this seems not to be the case for distinguishing algebraic growth from Dirac-coalescents: Extreme growth (large *γ*) seems to produce an almost star-shape like genealogy - consequently the distance to a Dirac coalescent with *ψ* close to 1 becomes very small (recall that *ψ* = 1 exactly corresponds to the star-shaped coalescent).

### Mean misclassification & posterior probabilities for the ABC approach

In this section, we analyse in how far an ABC approach using the nSFS (resp. the folded nSFS, abbreviated as nfSFS) and their lumped variants as summary statistic support our claim that one can distinguish between exponential growth `E` and the Beta(2−*α*, *α*)-coalescent `B` as well as between `E`, `B` and Dirac coalescents `D` or between `B`, `D` and algebraic growth `A`. The distinction ability of the ABC model comparison is assessed based on the simulation procedure and notation described in the methods section. Priors were uniform over the full range of coalescent parameters in models `B`, `D`, and uniform until a maximal cutoff for `E` (on [0, *β*_max_]) and `A` (on [0, *γ*_max_]). We discretized the parameter range for the growth models by using increments of 1 or 10 for exponential growth (the first used in all multiple-model comparisons, the latter in the pairwise comparisons between `E` and `B`) and increments of 1 for algebraic growth. If not specified otherwise, we used *β*_max_ = *γ*_max_ = 1000. We fixed a sample size *n* = 200. The number of replications was set to *n*_r_ = 2 × 10^5^. See Tables 1-3 and S4-S8 for the estimates of posterior probabilities and misclassification probabilities (some with one replication) with various degrees of lumping and various parameter settings for *n* = 200. For an example with higher sample size *n* = 1278, see Table S9.

**Table 1:**
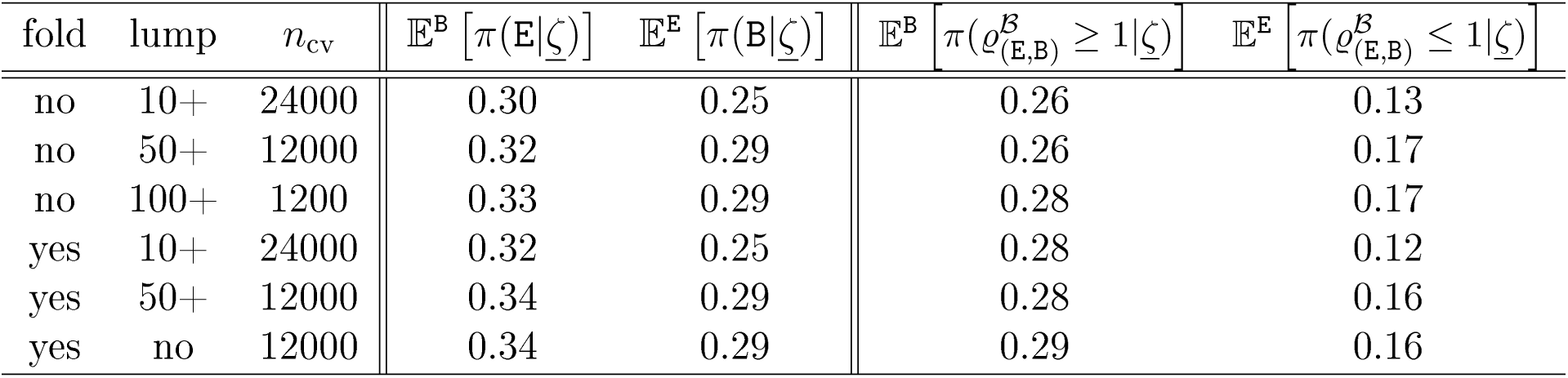
Approximations of the mean posterior probabilities and misclassification probabilities for the ABC model comparison between models `E` and `B` for tolerance *x* = 0.01, *s* = 15 segregating sites; and using either the nSFS or the nfSFS as summary statistics. *n*_cv_ denotes the number of cross-validations; ‘lump’ indicates which mutation classes are lumped into one class.

**Table 2:** Approximations of the mean posterior probabilities for the ABC model comparison between models `E`, `B`, `D` for tolerance *x* = 0.005, sample size *n* = 200 and *s* = 15 or *s* = 75. The nfSFS was used as summary statistics.

| $s$ | lump | $n_{\text{cv}}$ | $\mathbb{E}^{\mathbf{B}} [\pi(\mathbf{E}\vert \underline{\zeta})]$ | $\mathbb{E}^{\mathbf{B}} [\pi(\mathbf{D}\vert \underline{\zeta})]$ | $\mathbb{E}^{\mathbf{E}} [\pi(\mathbf{B}\vert \underline{\zeta})]$ | $\mathbb{E}^{\mathbf{E}} [\pi(\mathbf{D}\vert \underline{\zeta})]$ | $\mathbb{E}^{\mathbf{D}} [\pi(\mathbf{B}\vert \underline{\zeta})]$ | $\mathbb{E}^{\mathbf{D}} [\pi(\mathbf{E}\vert \underline{\zeta})]$ |
| --- | --- | --- | --- | --- | --- | --- | --- | --- |
| 15 | 10+ | 24000 | 0.29 | 0.12 | 0.24 | 0.02 | 0.55 | 0.03 |
| 15 | 50+ | 12000 | 0.39 | 0.10 | 0.22 | 0.02 | 0.54 | 0.05 |
| 15 | no | 12000 | 0.42 | 0.10 | 0.21 | 0.02 | 0.55 | 0.07 |
| 75 | 10+ | 24000 | 0.22 | 0.09 | 0.12 | 0.00 | 0.11 | 0.01 |
| 75 | 50+ | 12000 | 0.26 | 0.10 | 0.12 | 0.00 | 0.11 | 0.01 |

**Table 3:** Approximations of the mean posterior probabilities for the ABC model comparison between models `A`, `B`, `D` for tolerance *x* = 0.005, sample size *n* = 200 and *s* = 15 or *s* = 75. The nfSFS was used as summary statistics.

| $s$ | lump | $n_{cv}$ | $\mathbb{E}^B [\pi(A\vert \zeta)]$ | $\mathbb{E}^B [\pi(D\vert \zeta)]$ | $\mathbb{E}^A [\pi(B\vert \zeta)]$ | $\mathbb{E}^A [\pi(D\vert \zeta)]$ | $\mathbb{E}^D [\pi(B\vert \zeta)]$ | $\mathbb{E}^D [\pi(A\vert \zeta)]$ |
| --- | --- | --- | --- | --- | --- | --- | --- | --- |
| 15 | 10+ | 24000 | 0.02 | 0.12 | 0.01 | 0.03 | 0.16 | 0.55 |
| 15 | 50+ | 12000 | 0.02 | 0.11 | 0.01 | 0.03 | 0.24 | 0.57 |
| 75 | 10+ | 24000 | 0.01 | 0.09 | 0.01 | 0.05 | 0.10 | 0.29 |
| 75 | 50+ | 12000 | 0.01 | 0.08 | 0.01 | 0.05 | 0.15 | 0.32 |

The estimated error probabilities range from moderate to low values. Mean posterior probabilities 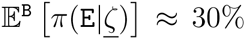 indicate a correct classification probability 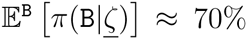, which shows that our method has good distinguishing ability. As expected, lower tolerance generally leads to smaller errors, as do larger mutation rates, while using the folded nSFS increases them. Appropriate lumping seems to decrease the error probabilities in many occasions, see e.g. Table 1 where a positive effect for strong lumping is observed for *s* = 15 segregating sites, whereas for *s* = 75 in Table S4, moderate lumping seems to be more appropriate (both tables show the comparison of models `B` and `E`). Not surprisingly exponential growth rates closer to zero are harder to distinguish from the Beta(2−*α*, *α*)-coalescent models than higher growth rates (see Tables S4 and S6). The ABC model comparison distinguishes especially well, even for *s* = 15, between exponential growth from Dirac coalescents and also algebraic growth from Beta(2 − *α*, *α*)-coalescent(Tables 3, S8, 2, S7). It should be noted that for relatively low number of segregating sites (*s* = 15), some comparisons (algebraic growth with Dirac coalescents, Beta coalescents with Dirac coalescents) can lead to common misclassification, but this effect vanishes for larger *s*. For *s* = 75, Dirac coalescents can be distinguished relatively well from Beta(2 − *α*,*α*)-coalescent (Tables 3, S8, 2, S7).

## Discussion

The development of methods to distinguish between different (time-changed) coalescent scenarios for the underlying genealogy of a population on the basis of observed data is an important task, in particular since the choice of an underlying coalescent model affects the estimated coalescent mutation rate 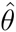 via (3) and a potential real-time embedding of the genealogy based on (5). The identification of an appropriate coalescent model may also give hints about the underlying reproductive mechanisms present in a population. By way of example, multiple-merger coalescents may indicate presence of HFSOD in the population.

While inference methods for distinguishing population growth from the usual Kingman coalescent have been extensively studied, see e.g. Tajima (1989a), Slatkin and Hudson (1991), Rogers and Harpending (1992), Kaj and Krone (2003) and Sano and Tachida (2005), and sophisticated theoretical results on the question of identifiabilitv of demographic histories have been obtained (cf. eg. Myers *et al.* (2008), Bhaskar and Song (2014) and Kim *et al.* (2014)), none of these studies have addressed multiple merger coalescents. In fact, only few results, eg. on the statistical properties of the SFS in multiple merger coalescents, see Birkner *et al.* (2013b), are available.

For the particular case of distinguishing multiple merger coalescents from population growth scenarios, this decision problem is complicated by the fact that the patterns of genetic variation produced by the two demographic effects, and summarized in the SFS, are expected to be similar: Both lead to an excess of singletons compared to a classical Kingman-coalescent based genealogy. However, while it is usually possible to match the predicted number of singletons with the observed number in various special cases for both models, the bulk and tail of the spectrum will typically differ, cf. Figure 1 for some examples.

The present paper is thus aimed at exploiting and quantifying these differences. However, for feasibility, we had to restrict both the scope and the employed methods of our analysis. The first (restrictive) decision in the design of our analysis was the selection of certain subfamilies of Lambda-coalescents and demographic growth scenarios that we deemed suitable for investigation. The reason for restricting to sub-classes of Lambda-coalescents is that the full class of multiple merger coalescents is in one-to-one relation with the uncountable and non-parametrie set of finite measures Λ on [0, 1], which drastically complicates statistical questions, while most of these coalescents do not appear to have a clear biological motivation in terms of a natural underlying population model. Considering the whole Lambda-coalescent class would also raise theoretical questions related to the unique identifyability of multiple merger coalescents on the basis of the SFS, related Myers *et al.* (2008) and Bhaskar and Song (2014), which is mathematically challenging and outside the scope of this work.

Hence, in case of the multiple merger coalescents, we restricted our attention to the class of Beta-coalescents (`B`) and the Dirac-coalescents (`D`). These classes appeared particularly interesting to us, since they both interpolate in a parametric way between the boundary points of Kingman-coalescent (`K`) via the Bolthausen-Sznitman coalescent (`B` at *α* = 1) to the ‘star-shaped-coalescent’ (`B`, *α* = 0; `D`, *ψ* = 1) - where the whole genealogy collapses to a single line in a single large merger event - among the multiple merger coalescents, Beta-coalescents have been studied frequently in the literature (see, eg., Birkner *et al.* (2005), Bertoin and Le Gall (2003), Hallatschek and Neher (2013), Steinrücken *et al* (2013), Birkner *et al.* (2013b)) and are related to a population model with high fecundity and skewed offspring distribution (HFSOD), Schweinsberg (2003), Dirac-coalescents have been chosen for their simplicity from a mathematical standpoint and have also been investigated in Eldon and Wakeley (2006). The parameter of the Dirac coalescent has a clear interpretation as the fraction of the population that is replaced in each single HFSOD reproductive event.

For similar reasons, we restricted demographic scenarios to two basic parametric growth models. Exponential growth (`E`) is certainly a natural model in the presence of a supercritical branching population model without geographic or resource restrictions. Our second choice, algebraic growth model (`A`) appears maybe less natural, but can reflect situations in which there are spatial- or resource limitations and has been analysed in the mathematical literature, eg, Schweinsberg (2010). We refrain from more complicated scenarios, such as models with different epochs of exponential growth, and recent models including superexponential growth (Reppell *et al.*, 2014), which indeed could be investigated with similar methods.

Regarding statistical methodology, one could construct likelihood-based tests on the full two-dimensional parameter spaces for Π and *θ*, given by Θ^E^ and Θ^B^ as outlined in the methods section, however, this will likely yield considerable computational challenges. Instead, we opted to employ approximate likelihood-methods based on the ‘fixed-*s*-method’ as eg. done by Ramos-Onsins and Rozas (2002), reducing our test to a one-dimensional situation, where (14) does not depend on *θ* at all.

Based on this method, we derive an approximate likelihood-ratio test based on a Poissonisation of the SFS via (12) for interval hypotheses, including large ranges of parameters such as the growth parameter *β* in model (`E`) or the coalescent parameter *α* in mode (`B`). By considering the power of our test, a key result in this setup is that, even for moderate sample size, (`B`) and (`E`) can be distinguished reasonably well for substantial parts of the parameter space of *α* and *β*.

A well-known criticism of this method is its sensitivity upon the true yet unknown eoalescent mutation rate *θ*, cf. Markovtsova *et al.* (2001). We checked by rejection sampling (cf. the supplementary material), conditioning on *S* = *s*, that for various fixed values of (Π, *θ*) the rejection probability in our proposed test would be reasonably close to the ‘true’ rejection probability as long as the ‘true’ *θ* is close enough to the Watterson estimate 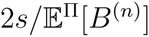, in line with similar observations (for different test statistics) made by Wall and Hudson (2001).

Given additional information about the exact coalescent and growth parameters could lead one to test the point hypotheses (eg. (`B`) with fixed *α* vs. (`E`) with fixed *β*). Indeed, in this case, higher power can be achieved, even for relatively small numbers of segregating sites (*s* = 20), as expected (data not shown).

Distance plots (eg. Figure 3) over two-dimensional parameter ranges indicate a onedimensional curve along which the minimal distance is reached. Note that both approaches (maximum-(approximate)-likelihood and minimum *ℓ*_2_-distance) could be linked if asymptotic normality of our estimators could be established - this is a theoretical question for future work.

Finally, we consider decision rules for the normalized spectrum 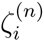 associated with models `A`, `B`, `E`, `D` based on a simple rejection-based ABC analysis. More sophisticated techniques are available (see Beaumont (2010) for an overview) that may improve the prediction accuracy. Empirical misclassification probabilities show, for a reasonable sample size of *n* = 200, at least moderate success in distinguishing between the 4 model classes even for as few as *s* = 15 segregating sites. Note though that depending on the model class comparison to perform, reasonable error probabilities may only be achieved at higher mutation rates/a higher number *s*. This indicates that the genealogies produced by the different model classes (at least for suitable sample sizes) are different enough to be distinguished, but that mutation rates have to be high enough so that these differences are mirrored in the SFS.

In practice, our results could be used to design studies that allow to distinguish between different conjectured scenarios with suitable power. For example, in marine species, such as Atlantic cod (cf. eg., Birkner *et al.,* 2013b) and Pacific oysters (Sargsyan and Wakeley, 2008), it has been suggested that certain multiple merger coalescents could be more appropriate to describe underlying genealogies, and a reproductive mechansism (HFSOD) for population models has been proposed. We have put this to a test by performing an ABC model comparison between our four model classes for the Atlantic cod data of Árnason (2004). The model comparison clearly rejected both Dirac coalescents and algebraic growth as potential models.

While our ABC analysis indicates exponential growth is slightly favored over the Beta (2 − *α*, *α*)-coalescent, evidence is not really strong enough to rule out the latter model class. This may indicate that the SFS information of the Árnason (2004) data does not have enough polymorphic sites to distinguish between `E` and `B` clearly enough, our posterior predictive checks revealed that likely neither model class explains the data completely. However, rejection of the `D` and `A` model classes suggest that models that predict star-shaped genealogies don’t fit the data well. Árnason (2004) uses a maximum likelihood estimation method (Kuhner *et al.*, 1998) and standard tests of neutrality (Tajima, 1989b; Fu, 1997) to rule out exponential population growth.

At this point we would like to point out that while our methods and results are exemplified in certain special coalescent and growth models, they could be modified to cover different frameworks.

Before ending the discussion we wish to comment on a few interesting side-issues that appeared during the analysis.

*Non-monotonicity of the power function.* At first glance, the observed non-monotonicity of the power function in the exponential growth parameter *β* when comparing with certain multiple merger coalescents (cf. e.g. Figures 2) may appear strange. However, the following example may suggest a heuristic way to understand such behaviour in a relatively simple special case. Suppose one wants to distinguish between an exponential growth model and a multiple merger coalescent with a substantial Kingman component and a small weight on large multiple mergers (eg. as in the Dirac-coalescent with *ψ* close to 1). That means that most of the time, the multiple merger coalescent will behave like a Kingman-coalescent (producing frequent binary mergers), but with small rate, comprehensive multiple mergers may occur. Certainly, when the growth parameter *β* is small, the exponential growth model will yield a pattern of variability close to a Kingman coalescent, and hence the power of a test to distinguish between both will be small if the Kingman component has weight close to 1. As *β* increases, the power to distinguish from a Kingman coalescent will increase, in line with intuition. However, as *β* becomes very large, lineages will coalesce after *very* short time in the exponential growth model. Such a scenario is certainly different from a Kingman coalescent, but could produce patterns of variability closer to a multiple merger coalescent with a drastic merger after very short time. Seeing such a merger in the very recent past has some cost (according to the weight Dirac component near 1), but appears more likely than observing large amounts of Kingman-like mergers within a unnaturally short time interval, thus leading to a relative decrease in power of associated test. This last effect is nicely illustrated by the upper-right scenario in Figure 4 in the case of large (algebraic) growth parameter.

*Effect of ‘lumping’.* It is intriguing to see that using the complete nSFS as summary statistics in the ABC approach can yield higher errors than using intermediate (resp. strong) lumpings of the nSFS. A possible explanation is as follows. Consider the approximate likelihood function (12). Assume that the distribution of the SFS is approximately composed of independent Poisson distributions with parameter 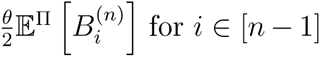. For a Poisson-distributed random variable *X* with parameter *κ*, we have 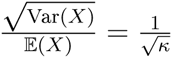, thus showing that smaller Poisson parameters yield a higher amount of variation relative to their expected value. Hence, classes in the SFS with small underlying branch lengths (which tend to be in the right tail of the SFS) and/or a low mutation rate show relatively more variation compared to their contribution to the total number of mutations than those with longer branches or if the mutation rate is higher. Lumping such classes together, under (12), yields again a Poisson-distributed lumped class, but with Poisson parameter being the sum of parameters from the classes lumped together. Thus, the variation within this class relative to its contribution to the total number of mutations is reduced by lumping. If different coalescent models show different mean behaviour of (lumped) classes, lumping reduces noise and thus increases the chance to correctly identify the underlying model. Naturally, this effect is weakened by higher mutation rates and/or higher sample size *n* (e.g., consider the limit results for the SFS in Berestycki *et al.* (2013) and Kersting and Stanciu (2013)).

Thus, using an appropriate weighing of the variables in the nSFS (resp, SFS) should improve the power to distinguish between model classes. It would also be a worthwile future study to see whether a one-dimensional summary of the SFS similar to Tajima’s D or Fay and Wu’s H, as described in Achaz (2009), could yield similar or even higher power to distinguish between the model classes than the complete (possibly reweighted) nSFS.

## Acknowledgements

J. Blath and B. Eldon were supported by Deutsche Forschungsgemeinschaft (DFG) grant BL 1105/3-1, and M. Birkner by DFG grant BI 1058/2-1, as parts of SPP Priority Programme 1590. F. Freund thanks Luca Ferretti and Guillaume Achaz (SMILE, Collège de France, Paris) for discussions about the site-frequency spectrum.

We would like to thank two anonymous referees whose insightful comments and constructive criticism helped to improve the presentation.

## Supporting Information

### Multiple merger-coalescents and the model classes K, B and D

Recall that a multiple merger-or Lambda-coalescent, formally introduced by Pitman (1999); Sagitov (1999), and Donnelly and Kurtz (1999), is a partition-valued exchangeable coalescent process determined by a finite measure Λ on [0, 1] which governs the dynamics of the process: If there are currently *b* blocks in the partition (i.e. *b* active ancestral lineages), *k* out of them merge at rate

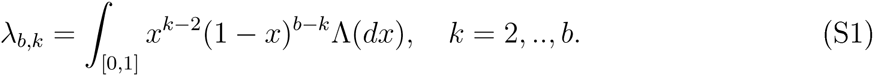

For an overview of the theory see e.g. Berestycki (2009) or, with a biological perspective, Tellier and Lemaire (2014). When Λ is associated with the beta-distribution with parameters 2 − *α* and *α* for 1 ≤ *α* < 2 (Schweinsberg, 2003), these rates can be given explicitly by

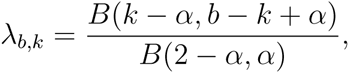

where *B*(·, ·) is the classical Beta-function. Such coalescents will be called Beta-coalescents, and constitute the model class `B`.

When Λ is associated with the Dirac coalescent (Eldon and Wakeley, 2006), that is, Λ(*dx*) = *δ*_{*ψ*}_(*dx*), for *ψ* ∈ [0, 1], we are in class `D`. Here, for *ψ* ∈ (0, 1], the rates are given by

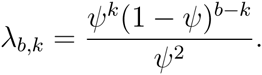

Both classes intersect in the Kingman coalescent (model `K`), which corresponds to *α* = 2 and *ψ* = 0, and of course has coalescence rates

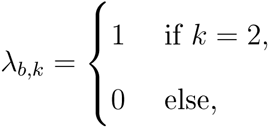

ie. only binary mergers are allowed. The Beta- and the Dirac coalescent each introduce a *coalescent* parameter (*α*, *ψ*), which can be estimated from genetic data (Eldon, 2011; Birkner *et al.*, 2013b; Birkner and Blath, 2008; Steinrücken *et al.*, 2013).

### Population models leading to coalescent classes K, B and D

It is well-known that the classical Wright-Fisher and the Moran model have scaling limits whose genealogy is described by a Kingman coalescent. For the more general Lambda-coalescents, Möhle and Sagitov (2001) give a full classification of all Cannings models that lead to any given Lambda-coalescent. The relevant time-scaling is determined by *c_N_*, the probability that in a population of size *N*, two distinct ancestral lineages merge in the previous generation. It is important to keep in mind that many different population models can lead to the same limiting coalescent, and also that the timescale, determined by *c_N_*, may vary between different models having the same limit. For the Kingman coalescent, the classical Wright Fisher model converges on the time-scale *c_N_* = 1/*N*, whereas for the Moran model, it is of order 1/*N*^2^.

A popular model that leads to the Beta(2 − *α*, *α*)-coalescent has been introduced by Schweinsberg (2003). For this model, the relevant time-scale is of order 1/*N^α^*^−1^. Here, single individuals can produce positive fractions of the next generation in a single reproductive event (an instance of ‘HFSOD’) that can be related to stable branching processes, cf. Birkner *et al.* (2005). The size of the reproductive event is random and governed by the Beta-distribution. For details we refer to Schweinsberg (2003), and for a discussion of its biological relevance eg. to Steinrücken *et al.* (2013).

The Dirac coalescent has been investigated in Eldon and Wakeley (2006). It has a particularly simple interpretation: Given the coalescent parameter *ψ* ∈ (0, 1], in each ‘substantial’ reproductive event, a fraction of 100 · *ψ*% of the generation die and are replaced by the offspring of a single parent (there can be other, ‘non-substantial’ reproductive events which, though potentially frequent, become invisible in the limit). This is an extreme case of HFSOD, and biologically it seems difficult to justify why the fraction *ψ* should always be the same. However, it is mathematically simple and interpolates between the Kingman coalescent *ψ* = 0 and the star-shaped coalescent *ψ* = 1, thus we included it in our study. For details see Eldon and Wakeley (2006).

### Population with varying population size and the classes E and A

In Kaj and Krone (2003), a time-changed *n*-coalescent under a general model of variable population size is derived. More precisely, the authors consider a haploid Wright-Fisher model with population size *N* at generation *r* = 0 and consider a population size process *M_N_*(*r*), 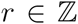 of the form *M_N_*(*r*) = *NX_N_*(*r*), 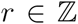, that is, *X_N_*(*r*) describes the ‘relative population size’ at generation *r*. Under the assumption that *X_N_*(⌊*Nt*⌋), 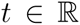 converges to something non-degenerate (ie. bounded away from 0 and ∞), they get the well-known limiting result that a time-changed Kingman coalescent describes the genealogy, where the infinitesimal coalescence rates are given by 1/*ν*(*s*), with
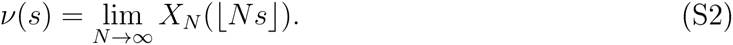

Our exponential growth model `E` corresponds to a Kingman-coalescent with exponentially growing coalescence rates *ν*(*s*) = *e^βs^*, for *β* ≥ 0, and can be obtained from a a growth rate of *β*/*N* per generation in the pre-limiting model, ie. *N_k_* = *N*(1 + *β*/*N*)*^k^*. Indeed,

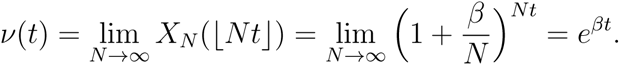

Thus, the size *Nt* generations ago is approximately *Ne*^−*βt*^.

The model class `A` is given by Kingman-coalescents with algebraically growing coalescence rates, ie. *ν*(*s*) = *s^γ^*, for *γ* ≥ 0. Note that if *γ* = 0 or *β* = 0, we recover the Kingman coalescent and are back in class `K`,

A population model for algebraic growth was considered in (Schweinsberg, 2010, Section 1.4): Fix a population size *N* at the present generation 0, and for notational convenience also for generation −1 (this short period of constant population size will become irrelevant after time-rescaling). For a fixed growth parameter *γ* > 0, the population size at the *k*-th generation before the present (for 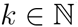) is assumed to be ⌈*Nk*^−*γ*^⌉. Measuring time in units of size 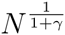 yields the limiting infinitesimal coalescence rate

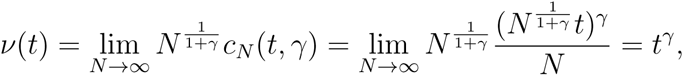

where *c_N_*(*t*, *γ*) is the probability that two individuals in generation 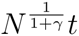 choose the same ancestor (uniformly out of the 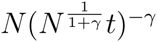 individuals alive in that generation). Consider the time-change (for the scaling limit as *N* → ∞)

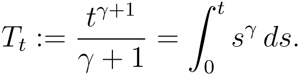

Then, the genealogy of the algebraic growth model at previous generation *t* equals in law the state of a classical Kingman coalescent at time *T_t_*. See Schweinsberg (2010) for details.

### The expected SFS under variable population size

The effect of fluctuations in population size on the SFS has been investigated in various articles, see eg. Griffiths and Tavaré (1998), who derive an analog of (1), and Kaj and Krone (2003) who link the Wright-Fisher approximation (with fluctuating population size) with the limiting genealogy.

Recursions for the expected values and covariances of the site-frequeney spectrum associated with moderate fluctuations in population size will now be briefly discussed. We will in particular consider numerically tractable recursions for the model classes `E` and `A`, based on work by Polanski *et al.* (2003) and Polanski and Kimmel (2003),

Consider a time-inhomogeneous Kingman coalescent, started in *n* lineages, where each pair of lines present at time *t* ≥ 0 merges at a rate *ν*(*t*). Then, the expected frequency spectrum 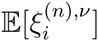, *i* ∈ [*n* − 1] is again of the form (1), and the time-change *ν* enters only in the distribution of the 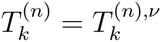, 2 ≤ *k* ≤ *n*, that is, the distribution of the lengths of the time intervals of the block-counting process 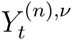 during which there are exactly *k* lineages.

To evaluate 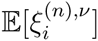 one needs information about 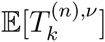. Define

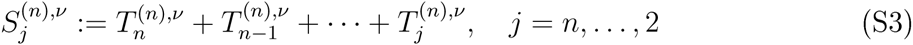

to be the time at which the block counting process *Y*^(*n*),*ν*^ jumps from *j* to *j* − 1 lineages (with the convention 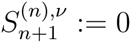). Abbreviate, for *t* ≥ 0 and *j* ∈ 2,…, *n*,

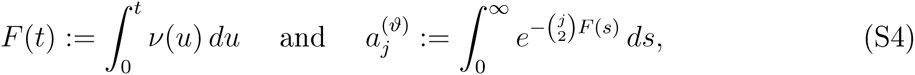

assuming that the first integral in (S4) is finite. It is possible to compute the marginal density of 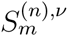 using the well-known fact that the density of a convolution of exponentials with different rates can be written as a linear combination of exponential densities,

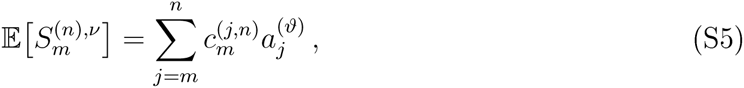

where

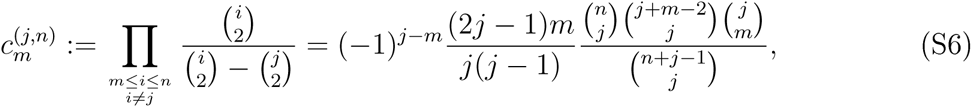

(put 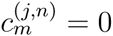 for *j* < *m*).

Polanski and Kimmel (2003) obtain numerically stable and efficient recursions to compute 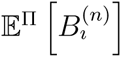 associated with any time-changed Kingman coalescent Π as follows. For *ϑ* denoting the growth parameter associated with process Π,

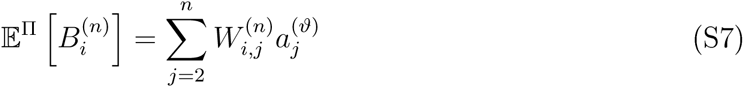

where the constants 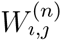 can be computed recursively (Polanski and Kimmel, 2003);

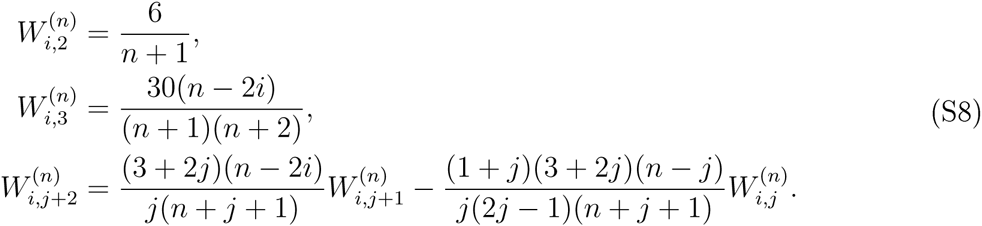

We now specify the main ingredient 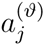 (depending on *F* (*t*), *t* ≥ 0 and hence *ν*(*t*), *t* ≥ 0) explicitly for two important special cases:

**a) Exponential growth.** In the case of an exponentially growing population with growth parameter *β* that is, *ν*(*t*) = *e^βt^*, we have

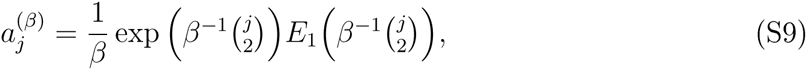

where

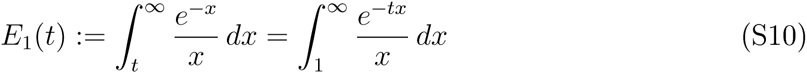

is an exponential integral function, c.f. e.g. (Abramowitz and Stegun, 1964, 5.1.1). One can use numerical integration schemes to compute *E*_1_(*t*) for smaller values of *t* (eg. *t* < 50). For larger values of *t*, one can use the approximation

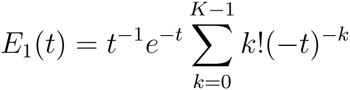

(Milgram, 1985), which has error of order *O* (*K*!*t* ^−*K*^).

**b) Algebraic (‘power law’) growth.** In the case of algebraic growth of the form *ν*(*t*) = *t^γ^* for some *γ* > 0, we have

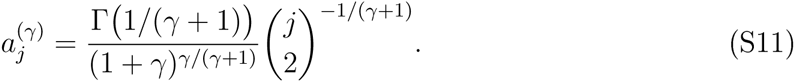

Based on Equation (23) in Fu (1995), it is also possible to compute the *variance* and the *covariances* of the SFS based on expressions for 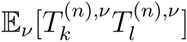, 2 ≤ *k*, *l* ≤ *n*, which in turn can be obtained from

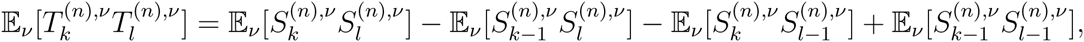

noting that, in the above notation,

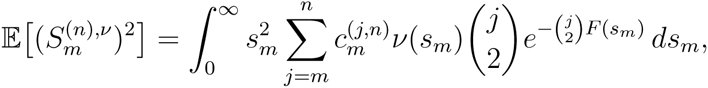

and

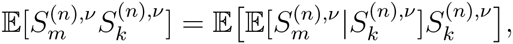

where 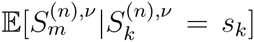 can be computed (it is the expectation under a regular conditional probability) as in (S5) replacing *ν* by 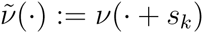, 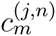 by 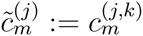 and *F* by 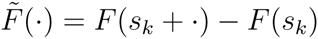.

### The estimate 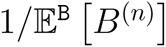 as a function of *α*

**Figure S1:**
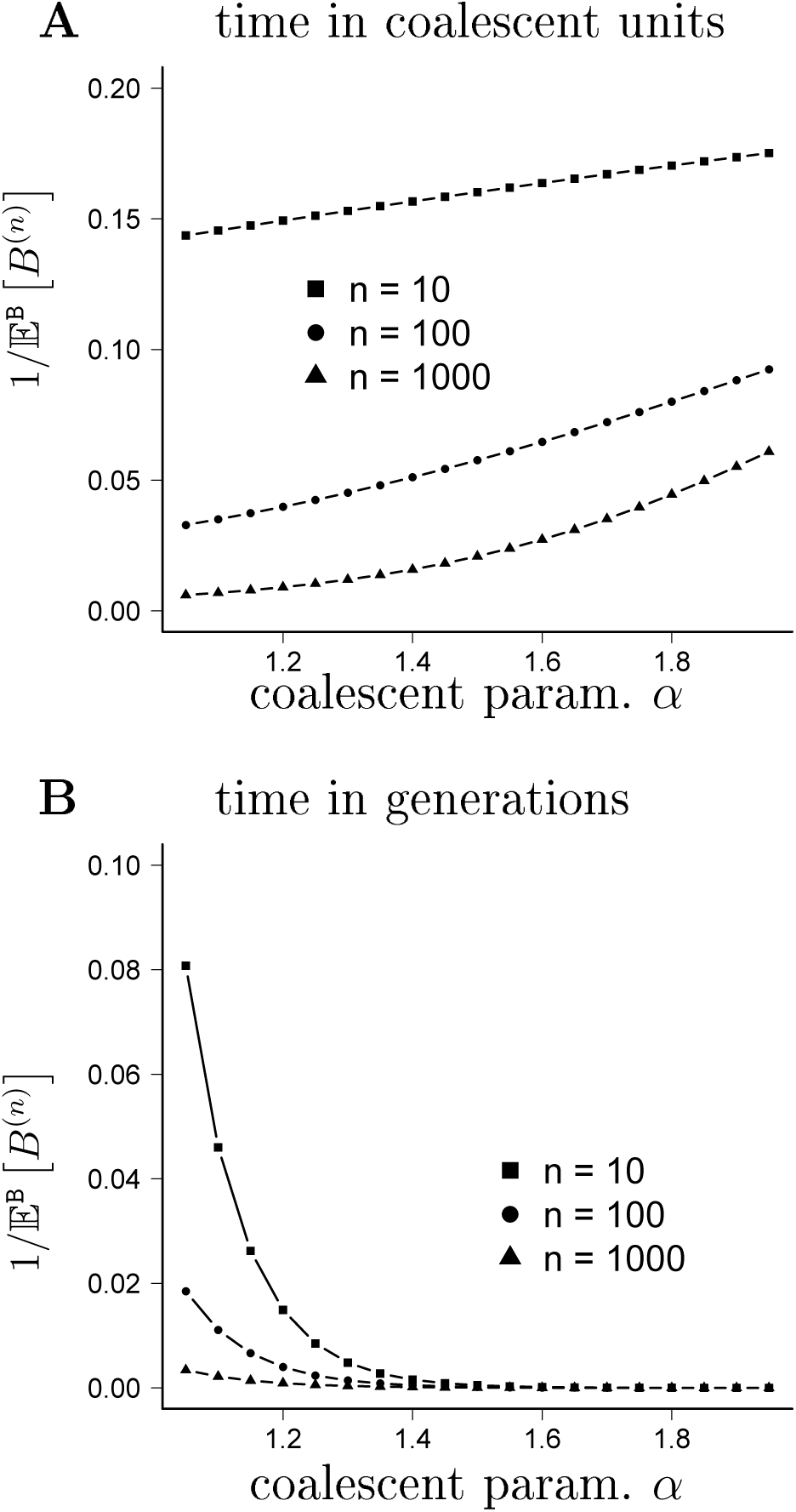
Graphs of 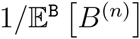, the estimated value of *θ*/2 per observed mutation when using the Watterson estimator (3) as a function of *α* (**A**), compare with (3); and the estimated value of *μ* per observed mutation (B), using (5) together with (3), and assuming the timescale *c_N_* = *N*^1−*α*^. The number of leaves *n* are as shown. In **B**, time is converted into generations by multiplying 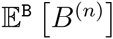 with *N^α^*^−1^ when *N* = 10^5^.

### Approximate insensitivity of the expected normalized SFS w.r.t. *θ*

In this section, we argue that for a random genealogical tree 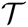 with *n* leaves whose law is governed by a given coalescent mechanism Π, the expected nSFS 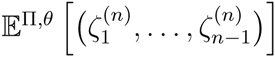 when the coalescent mutation rate is *θ* > 0 is approximately constant as a function of *θ*. This is useful because it in a sense allows to “factor out” (i.e, ignore) the mutation rate parameter from a test problem when comparing different Π’s. This also means that - at least when the observed number |*ξ*^(*n*)^| of segregating sites is reasonably large - the exact observed value |*ξ*^(*n*)^| does not add much additional information for tests based on the SFS.

Indeed, we can compute

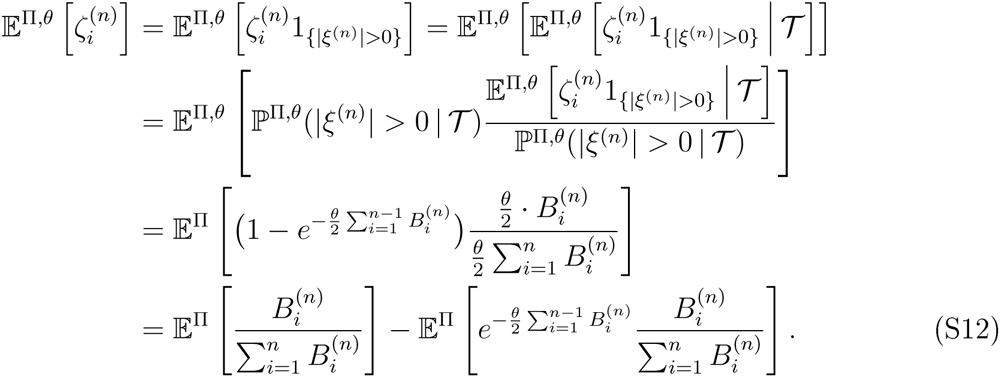

Here, 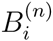 denotes the total length of all branches in 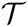 which subtend *i* leaves for *i* = 1, …, *n* − 1 and in the third line we used Lemma S1.1 below together with the fact that given 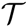 and *θ*, 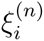, *i* = 1, …, *n* − 1 are independent and each 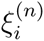 is Poisson distributed with mean 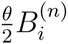. Note that the first term in (S12) is independent of *θ* and the “correction” term is small unless *θ* is very small or 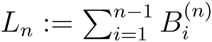, the total length of 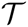, is small under Π with substantial probability. Note that for each of the coalescent processes we consider in this investigation, it does hold that *L_n_* → ∞ as *n* → ∞. Simulations also indicate that the distribution (not only the mean) of 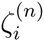 does not depend much on *θ* (data not shown).

#### Lemma S1.1.

*Let X*_1_, *X*_2_ *be independent Poisson-distributed variables with parameters a and b. Then*,

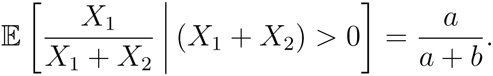

*Proof*. *X*_1_ +*X*_2_ as a sum of independent Poisson distributed random variables is again Poisson distributed with parameter *a* + *b*. We have

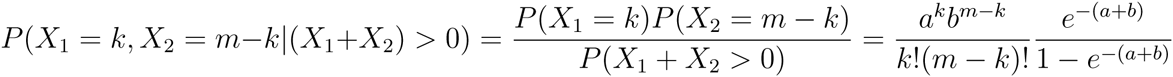

for 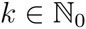, 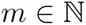 with *k* ≤ *m*. We compute

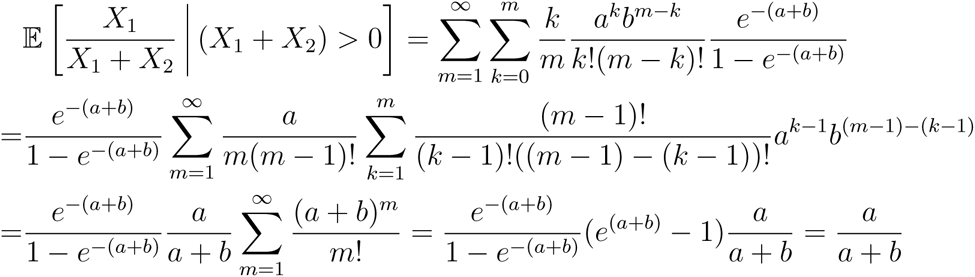

### Insensitivity of the fixed-*s*-method w.r.t. *θ*

To check the the robustness of our fixed-*s*-method against varying *θ* under rejection sampling (cf. e.g. Markovtsova *et al.* (2001), Wall and Hudson (2001)), we applied the following exact rejection sampling approach to simulate a coalescent tree conditional on a given number of observed segregating sites *s*. As input, the algorithm takes sample size *n*, number of segregating sites *s*, a coalescent model Π, and mutation rate *θ*, and returns a realisation of 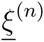 with 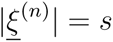.

**Rejection sampling algorithm :**

i. generate a coalescent tree according to Π, read off branch lengths 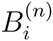,
ii. draw a total number of mutations *S* as realization of a Poisson random variable with 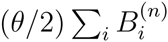,
iii. if *S* = *s* the required fixed number of segregating sites, keep the 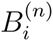, otherwise discard and draw again,
iv. throw uniformly *s* mutations on the tree, 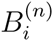, so that the probability of a mutation falling into class *i* is 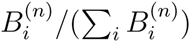.

We then computed (approximately via rejection-sampling) the size of a conditional distribution based test if one employs quantiles of the fixed-*s*-method derived from (8). Of course, the hope is that both are reasonably close to each other, and this seems to hold relatively well if *θ* is close to the Watterson estimate 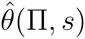 (9). In particular, the results (Tables (S1)–(S3)) show that the method is particularly robust against varying *θ* when exponential growth is taken as null model.

**Table S1:** Checking size of test given fixed-*s* quantiles associated with size *x*% and *α* ∈ {1, 1.5} with Beta(2 − *α*, *α*)-coalescent as null model, and exponential growth as alternative, using rejection sampling with mutation rate *θ* as shown. Sample size *n* = 100, segregating sites *s* = 50. The estimate (*θ_W_* (*α*)) is obtained from (9). All estimates from 10^5^ iterates.

| $\alpha$ | $x\%$ | $\theta$ ($\theta_W(\alpha)$) | size of test |
| --- | --- | --- | --- |
| 1.0 | 10% | 3.082453 ($\theta_W(1)$) | 0.10 |
|  |  | 2.0 | 0.13 |
|  |  | 3.0 | 0.10 |
|  |  | 5.0 | 0.07 |
|  |  | 7.0 | 0.06 |
| | 5% | 3.082453 ($\theta_W(1)$) | 0.05 |
|  |  | 2.0 | 0.07 |
|  |  | 5.0 | 0.03 |
|  |  | 7.0 | 0.02 |
| | 1% | 3.082453 ($\theta_W(1)$) | 0.01 |
|  |  | 2.0 | 0.02 |
|  |  | 5.0 | 0.01 |
|  |  | 7.0 | 0.002 |
| 1.5 | 10% | 5.7638 ($\theta_W(1.5)$) | 0.11 |
|  |  | 3.0 | 0.03 |
|  |  | 5.0 | 0.09 |
|  |  | 7.0 | 0.11 |
|  |  | 10.0 | 0.13 |
| | 5% | 5.7638 ($\theta_W(1.5)$) | 0.05 |
|  |  | 3.0 | 0.01 |
|  |  | 5.0 | 0.04 |
|  |  | 7.0 | 0.06 |
|  |  | 10.0 | 0.07 |
| | 1% | 5.7638 ($\theta_W(1.5)$) | 0.01 |
|  |  | 3.0 | 0.001 |
|  |  | 5.0 | 0.01 |
|  |  | 7.0 | 0.01 |
|  |  | 10.0 | 0.02 |

**Table S2:** Checking size of test given fixed-*s* quantiles associated with size *x*% as shown using rejection sampling with mutation rate *θ* as shown for exponential growth as null model, and Beta(2 − *α*, *α*)-coalescent as alternative. Sample size *n* = 50, segregating sites *s* = 25. The estimate (*θ_W_* (*β*)) is obtained from (9). All estimates from 10^5^ iterates.

| $\beta$ | $x\%$ | $\theta$ ($\theta_W(\beta)$) | test size |
| --- | --- | --- | --- |
| 1 | 10% | 7.895425 $\theta_W(1)$ | 0.10 |
|  |  | 5.0 | 0.10 |
|  |  | 7.0 | 0.10 |
|  |  | 9.0 | 0.10 |
|  |  | 11.0 | 0.10 |
| | 5% | 7.895425 $\theta_W(1)$ | 0.05 |
|  |  | 5.0 | 0.05 |
|  |  | 7.0 | 0.05 |
|  |  | 9.0 | 0.05 |
|  |  | 11.0 | 0.05 |
| | 1% | 7.895425 $\theta_W(1)$ | 0.01 |
|  |  | 5.0 | 0.01 |
|  |  | 7.0 | 0.01 |
|  |  | 9.0 | 0.01 |
|  |  | 11.0 | 0.01 |
| 10 | 10% | 16.33632 $\theta_W(10)$ | 0.10 |
|  |  | 12.0 | 0.13 |
|  |  | 14.0 | 0.12 |
|  |  | 18.0 | 0.10 |
|  |  | 20.0 | 0.10 |
| | 5% | 16.33632 $\theta_W(10)$ | 0.05 |
|  |  | 12.0 | 0.05 |
|  |  | 14.0 | 0.05 |
|  |  | 18.0 | 0.05 |
|  |  | 20.0 | 0.05 |
| | 1% | 16.33632 $\theta_W(10)$ | 0.01 |
|  |  | 12.0 | 0.01 |
|  |  | 14.0 | 0.01 |
|  |  | 18.0 | 0.01 |
|  |  | 20.0 | 0.01 |

**Table S3:**
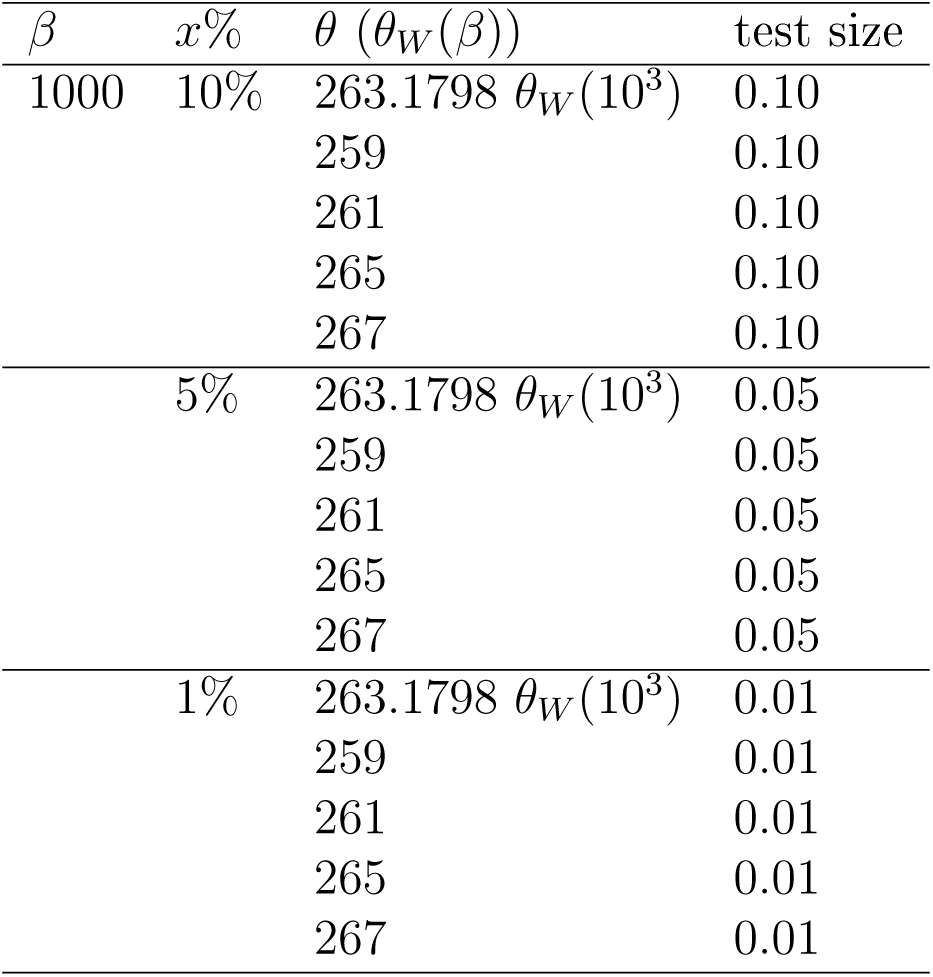
Checking size of test given fixed-*s* quantiles associated with size *x*% as shown using rejection sampling with mutation rate *θ* as shown for exponential growth as null model (*β* = 1000), and Beta(2 − *α*, *α*)-coalescent as alternative. The estimate (*θ_W_* (*β*)) is obtained from (9), Sample size *n* = 50, segregating sites *s* = 25. All estimates from 10^5^ iterates.

| $\beta$ | $x\%$ | $\theta$ ($\theta_W(\beta)$) | test size |
| --- | --- | --- | --- |
| 1000 | 10% | 263.1798 $\theta_W(10^3)$ | 0.10 |
|  |  | 259 | 0.10 |
|  |  | 261 | 0.10 |
|  |  | 265 | 0.10 |
|  |  | 267 | 0.10 |
| | 5% | 263.1798 $\theta_W(10^3)$ | 0.05 |
|  |  | 259 | 0.05 |
|  |  | 261 | 0.05 |
|  |  | 265 | 0.05 |
|  |  | 267 | 0.05 |
| | 1% | 263.1798 $\theta_W(10^3)$ | 0.01 |
|  |  | 259 | 0.01 |
|  |  | 261 | 0.01 |
|  |  | 265 | 0.01 |
|  |  | 267 | 0.01 |

### Estimation of power for 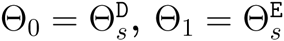

**Figure S2:**
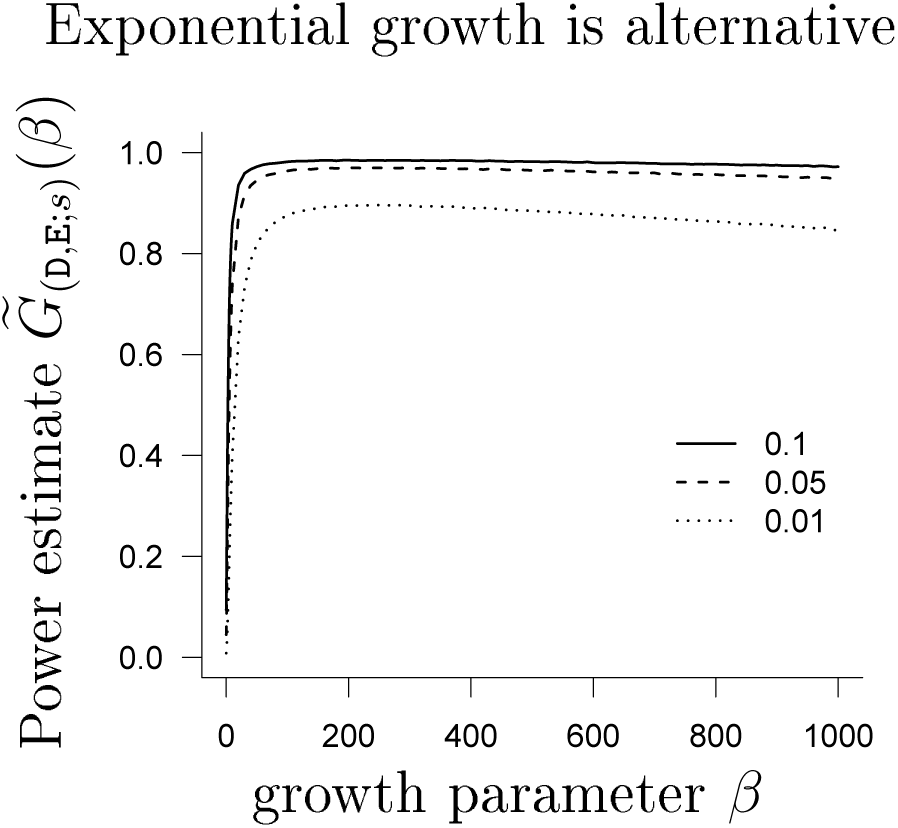
Estimate of 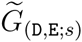 from (10) based on the approximate likelihood (12) as a function of *ψ* (no lumping) with number of leaves *n* = 100 and *s* = 50. The line types denote the size of the test as shown in the legend. The interval hypotheses are discretized to 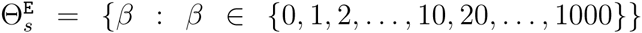 and 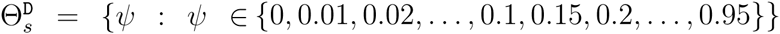. Reverting the hypotheses yield very similar results (not shown).

### Estimation of power for *s* = 300

**Figure S3:**
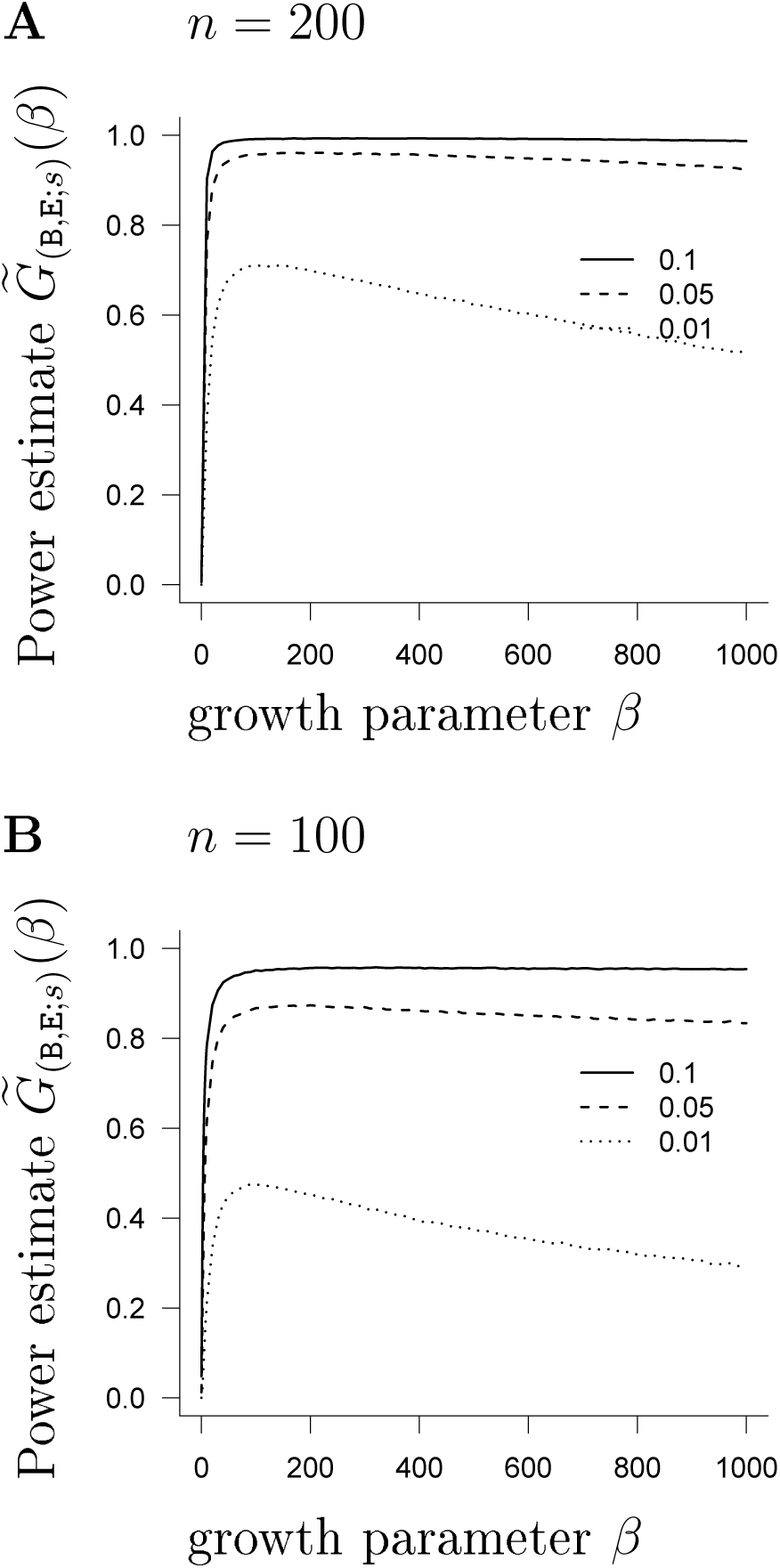
Estimate 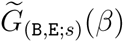 of power as a function of *β* for (**A**) *β* ∈ {0, 10,…, 1000}; (**B**) *β* ∈ {0, 1, 2,…, 9, 10, 20,…, 1000} when the Beta(2 − *α*, *α*)-coalescent is the null hypothesis, and the test statistic is 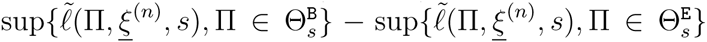 (7), with 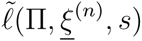 the log of the Poisson likelihood function (12) (no lumping). Values at *β* = 0 correspond to the Kingman coalescent. A total of 10^6^ replicates for both quantiles and power estimates.

**Figure S4:**
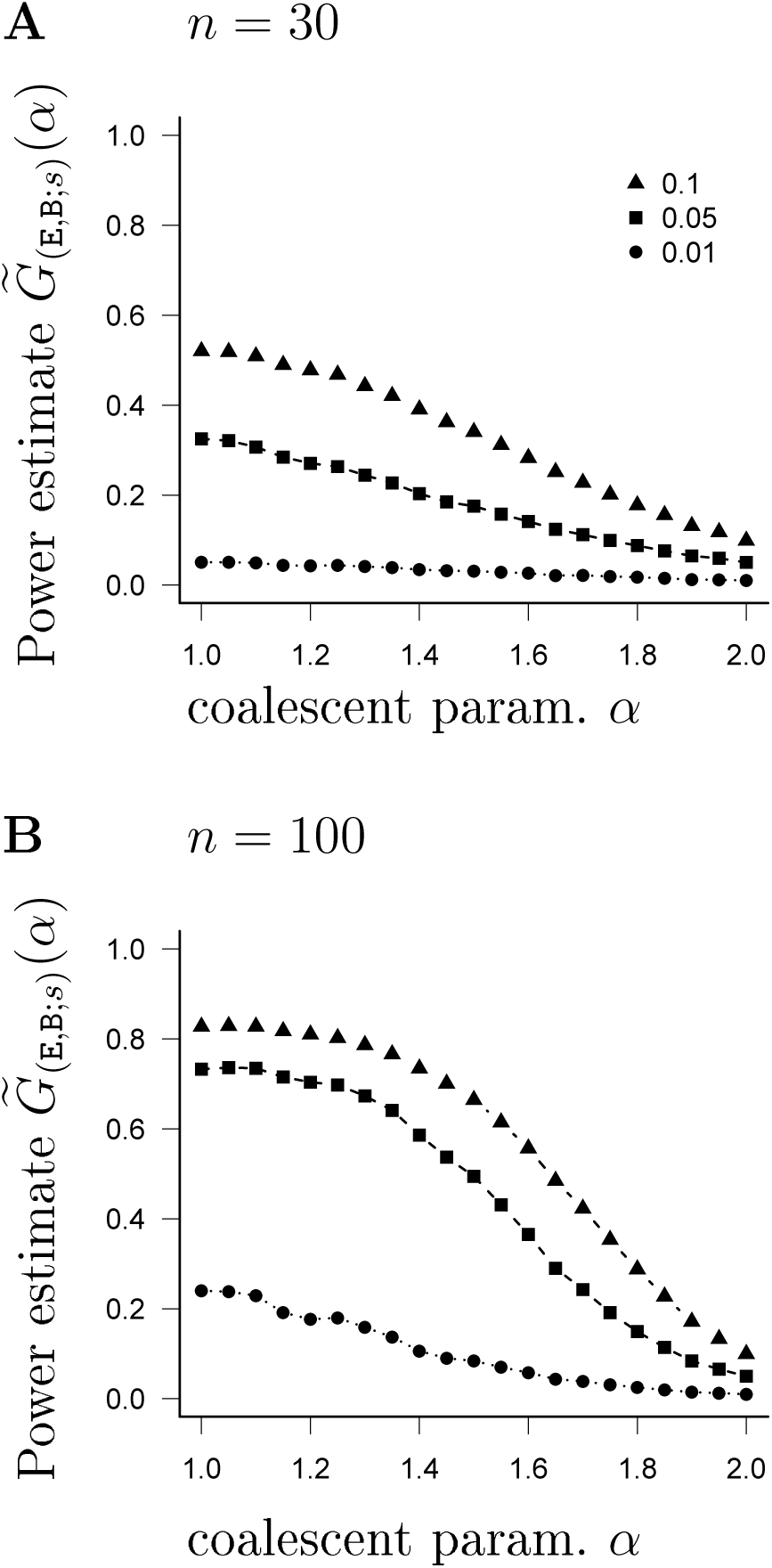
Estimate 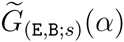 of power as a function of *α* for *α* ∈ [1, 2] when exponential growth (`E`) is the null hypothesis, Beta(2 − *α*, *α*)-coalescent (`B`) is the alternative, and the test statistic is 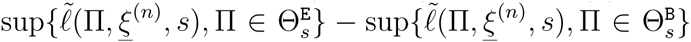 (7), with 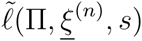 the log of the Poisson likelihood function (12) (no lumping). Values at *α* = 2 correspond to the Kingman coalescent; number of segregating sites *s* = 300; 10^6^ replicates for quantiles and power estimates.

**Figure S5:**
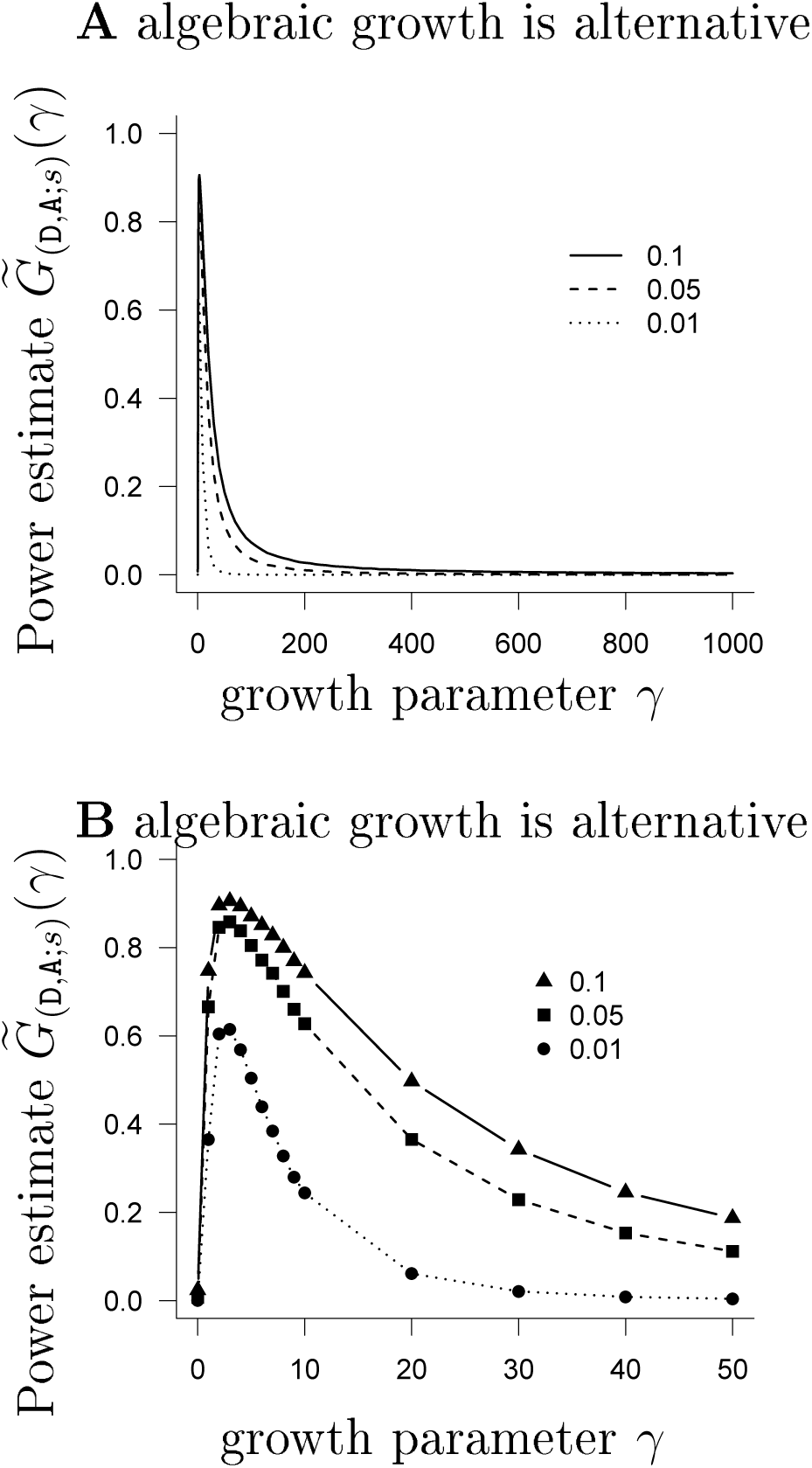
Estimate 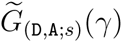 of power (10) between algebraic growth and the Dirac Lambda-coalescent when the test statistic is 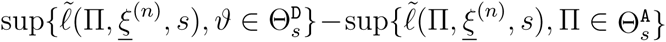, (7), with 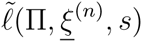 the log of the Poisson likelihood function (12) (no lumping); with *n* = 100 and number of segregating sites *s* = 50. The test sizes are as shown in the legend, The interval hypotheses are 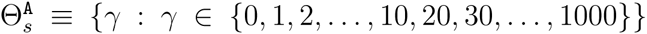 and 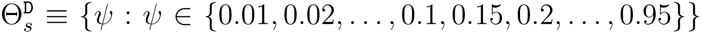. Values at *γ* = 0 correspond to the Kingman coalescent. Expected values were computed exactly, and quantiles and power estimated from 10^5^ replicates. Reverting the hypotheses shows a very similar pattern (results not shown). In **B**, we ‘zoom in’ on the range 0 ≤ *γ* ≤ 50.

**Figure S6:**
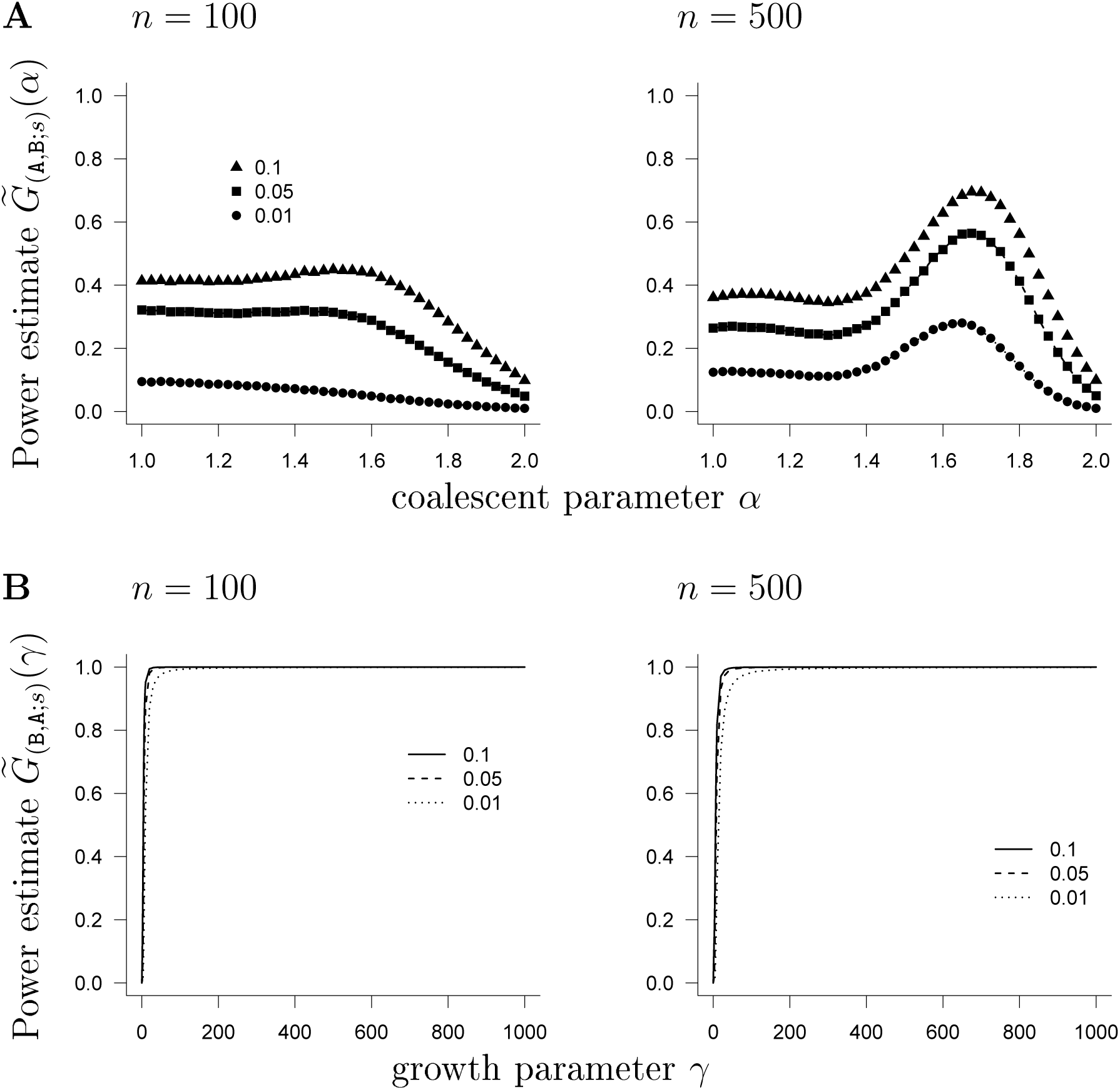
Estimate 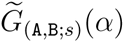 (**A**) and 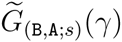 (**B**) of power (10) between algebraic growth and the Beta(2 − *α*, *α*)-coalescent when the test statistic is 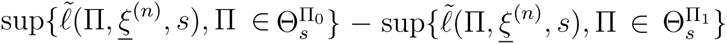 (7), with 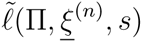 the log of the Poisson likelihood function (12) (no lumping); with number of leaves *n* as shown and number of segregating sites *s* = 50. The test sizes are as shown in the legend. The interval hypotheses are 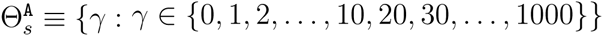 and 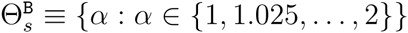. Values at *γ* = 0 and *α* = 2 correspond to the Kingman coalescent. Expected values were computed exactly, and quantiles and power estimated from 10^5^ replicates. In **A**, the Beta(2 − *α*, *α*)-coalescent is the alternative hypothesis; in **B**, algebraic growth is the alternative.

### Mean misclassification probabilities and posterior probabilities for ABC approach - alternative parameter choices

**Table S4:**
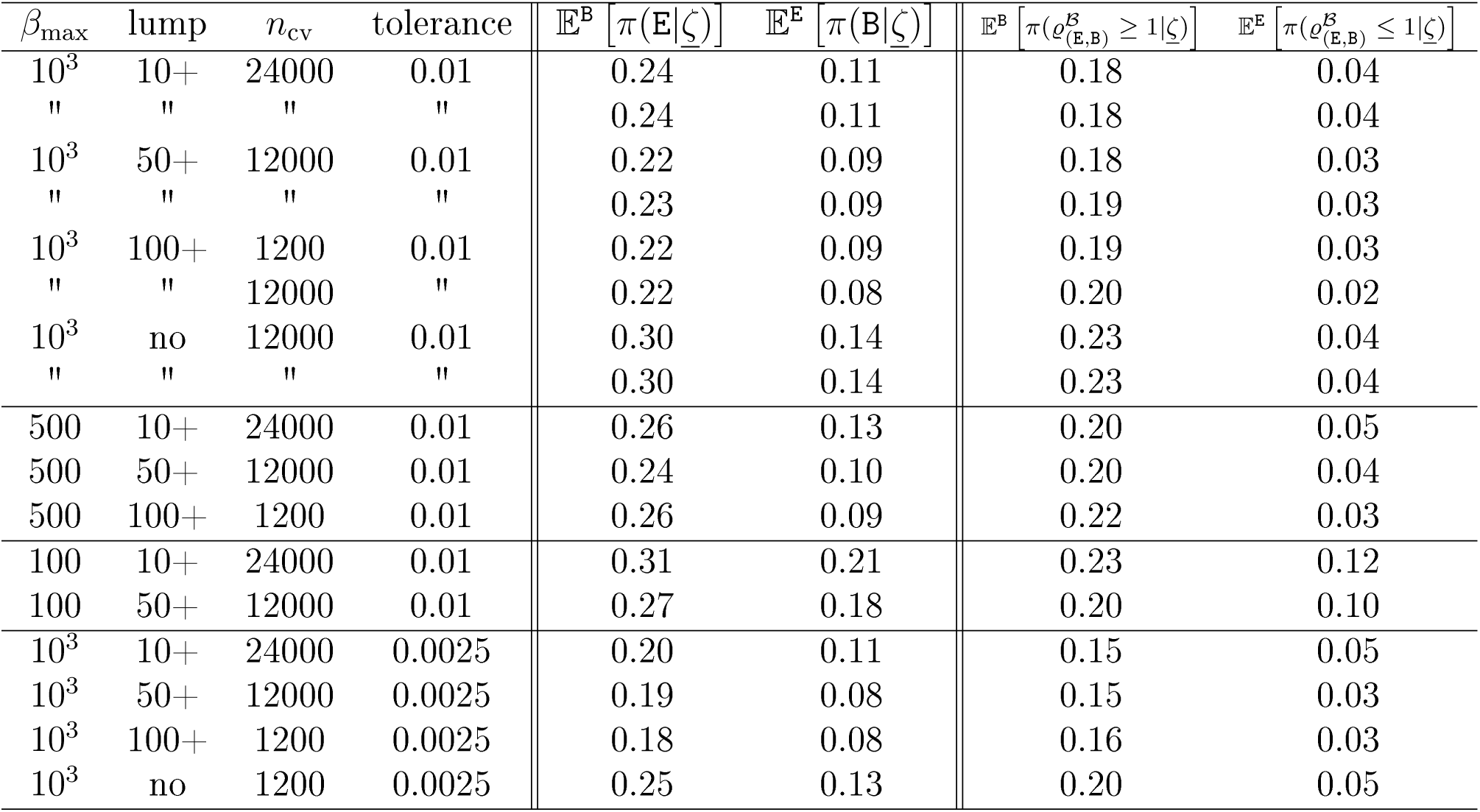
Approximations of mean posterior probabilities and misclassification probabilities for the ABC model comparison between `E` and `B` for different growth parameter ranges or tolerance rates. The nSFS is used as summary statistics. *β*_max_ denotes the maximal growth rate used in the growth model, *n*_cv_ denotes the number of cross-validations; ‘lump’ indicates which mutation classes are lumped into one class. An expected number *s* = 75 of mutations are assumed.

**Table S5:** Approximations of mean posterior probabilities and misclassifieation probabilities for the ABC model comparison between `E` and `B` for tolerance *x* = 0.0025 and sample size *n* = 200 and assumed expected number *s* = 15 of mutations. The nSFS is used as summary statistics. *n*_cv_ denotes the number of cross-validations ‘lumped’ indicates which mutation classes are lumped into one class.

| lump | $n_{cv}$ | $\mathbb{E}^B [\pi(E\vert \underline{\zeta})]$ | $\mathbb{E}^E [\pi(B\vert \underline{\zeta})]$ | $\mathbb{E}^B [\pi(\varrho_{(E,B)}^B \geq 1\vert \underline{\zeta})]$ | $\mathbb{E}^E [\pi(\varrho_{(E,B)}^B \leq 1\vert \underline{\zeta})]$ |
| --- | --- | --- | --- | --- | --- |
| 10 | 24000 | 0.28 | 0.24 | 0.23 | 0.14 |
| 50 | 12000 | 0.31 | 0.26 | 0.25 | 0.14 |
| 100 | 12000 | 0.33 | 0.27 | 0.28 | 0.15 |
| no | 12000 | 0.34 | 0.26 | 0.29 | 0.15 |

**Table S6:** Approximations of mean posterior probabilities and misclassification probabilities for the ABC model comparison between `E` and `B` for tolerance *x* = 0.001 and sample size *n* = 200, assumed expected number *s* = 15 of mutations and alternative prior ranges and distributions. The nSFS is used as summary statistics. *n*_cv_ denotes the number of cross-validations ‘lumped’ indicates which mutation classes are lumped into one class. For growth rate *β*, the prior is uniformly distributed on {*β*_min_, *β*_min_ + 10,…, *β*_max_}. For coalescent parameter *α*, the prior is uniformly distributed on [*α*_min_, *α*_max_]

| lump | $n_{cv}$ | $\beta_{\min}, \beta_{\max}$ | $\alpha_{\min}, \alpha_{\max}$ | $\mathbb{E}^B [\pi(E\vert \underline{\zeta})]$ | $\mathbb{E}^E [\pi(B\vert \underline{\zeta})]$ | $\mathbb{E}^B [\pi(\varrho_{(E,B)}^B \geq 1\vert \underline{\zeta})]$ | $\mathbb{E}^E [\pi(\varrho_{(E,B)}^B \leq 1\vert \underline{\zeta})]$ |
| --- | --- | --- | --- | --- | --- | --- | --- |
| 10 | 24000 | 0,100 | 1.5,2 | 0.39 | 0.34 | 0.30 | 0.23 |
| 50 | 12000 | 0,100 | 1.5,2 | 0.38 | 0.31 | 0.31 | 0.18 |
| 10 | 24000 | 100,1000 | 1,1.5 | 0.33 | 0.28 | 0.29 | 0.14 |
| 50 | 12000 | 100,1000 | 1,1.5 | 0.36 | 0.32 | 0.31 | 0.18 |

**Table S7:** Approximations of the misclassification probabilities for the ABC model comparison between models `E`, `B`, `D` for tolerance *x* = 0.005, sample size *n* = 200 and *s* ∈ {15, 75}. The folded nSFS was used as summary statistics. We use the abbreviation 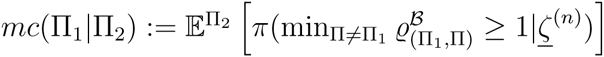, Π_1_, Π_2_ ∈ {`E`, `B`, `D`}.

| s | lump | $n_{cv}$ | $mc(E\vert B)$ | $mc(D\vert B)$ | $mc(B\vert E)$ | $mc(D\vert E)$ | $mc(B\vert D)$ | $mc(E\vert D)$ |
| --- | --- | --- | --- | --- | --- | --- | --- | --- |
| 15 | 10+ | 24000 | 0.27 | 0.07 | 0.12 | 0.01 | 0.62 | 0.01 |
| 15 | 50+ | 12000 | 0.39 | 0.06 | 0.08 | 0.01 | 0.60 | 0.03 |
| 15 | no | 12000 | 0.42 | 0.07 | 0.08 | 0.01 | 0.64 | 0.04 |
| 75 | 10+ | 24000 | 0.19 | 0.04 | 0.05 | 0.00 | 0.09 | 0.00 |
| 75 | 50+ | 12000 | 0.24 | 0.04 | 0.04 | 0.00 | 0.09 | 0.00 |

**Table S8:** Approximations of the misclassification probabilities for the ABC model comparison between models `A` `B` `D` for tolerance *x* = 0.005, sample size *n* = 200 and *s* ∈ {15, 75}. The folded nSFS was used as summary statistics. We use the abbreviation 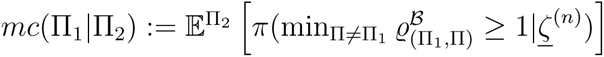, Π_1_, Π_2_ ∈ {`A`, `B`, `D`}.

| s | lump | $n_{cv}$ | $mc(A\vert B)$ | $mc(D\vert B)$ | $mc(B\vert A)$ | $mc(D\vert A)$ | $mc(B\vert D)$ | $mc(A\vert D)$ |
| --- | --- | --- | --- | --- | --- | --- | --- | --- |
| 15 | 10+ | 24000 | 0.01 | 0.06 | 0.01 | 0.04 | 0.15 | 0.53 |
| 15 | 50+ | 12000 | 0.01 | 0.06 | 0.01 | 0.04 | 0.18 | 0.52 |
| 75 | 10+ | 24000 | 0.00 | 0.03 | 0.01 | 0.06 | 0.09 | 0.25 |
| 75 | 50+ | 12000 | 0.00 | 0.03 | 0.01 | 0.05 | 0.14 | 0.27 |

### ABC analysis of the cytochrome *b* mtDNA data of Árnason (2004)

To investigate which model class (exponential growth `E`, algebraic growth `A`, Beta(2 − *α*, *α*)-coalescents `B`, Dirac coalescents `D`) fits better to the data, we use the ABC model comparison approach given the (lumped) nfSFS of the observed mitochondrial locus. The exponential growth model class is specified by an uniform prior for growth parameter *β* on {0, 1, 2,…, 1000}, the algebraic growth class by an uniform prior for growth parameter γ on {0,1, 2,…, 1000}, The class of Beta *n*-coalescents is specified by an uniform prior on {1, 1.01,…, 2} for the coalescent parameter *α*, the class of Dirac coalescents by an uniform prior on {0.01, 0.02,… 0.99} for the coalescent parameter *ψ* (we omit the star-shaped coalescent *ψ* = 1 because the observed SFS has not only singleton mutations, thus directly violating this model). We used two tolerance levels of 0.005 and 0.00125 and perform *n*_r_ = 200,000 simulations for each model class. See Table S9 for the approximated Bayes factors 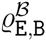 for the model comparison of the growth model and the Beta *n*-coalescent model using different lumps of the nfSFS as summary statistics. The Bayes factors 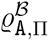, 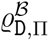 for Π ∈ {`E`, `B`} have maximal values of ≈ 0.01, 0.001 under all lumpings and both tolerances. The observed data fits slightly better to the growth model than to the Beta coalescent class, but not so much better that we could discard the Beta *n*-coalescents as possible genealogy models for this locus. The latter point is also highlighted by results for an ABC model comparison between only model classes `E` and `B` where all lumpings but 100+ again (slightly) favour the growth model, but for 100+ lumping this is reversed (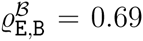 for tolerance 0.005). The Dirac coalescents and the algebraic growth model show neglectible support for all lumpings and thus we discard them as potential models.

**Table S9:** Approximated Bayes factor 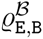 given the Atlantic cod mtDNA data

| lumping number | 10+ | 50+ | 100+ | 200+ | no |
| --- | --- | --- | --- | --- | --- |
| tolerance 0.005 | 7.79 | 2.23 | 2.09 | 2.97 | 2.98 |
| tolerance 0.00125 | 10.35 | 2.97 | 2.23 | 6.87 | 7.13 |

Jeffreys (1961) suggested interpreting Bayes factors according to the log_10_ scale. Lumping at 10 (Table S9) then gives at least ‘substantial’ 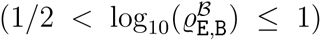 evidence against the Beta(2 − *α*, *α*)-coalescent in favor of exponential growth. Using Kass and Raftery (1995) suggestion of considering Bayes factors on 2log*_e_* scale gives ‘positive’ 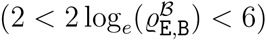 evidence in favor of exponential growth, based on lumping at 10.

Additionally to the ABC model comparison, we also evaluate which parameters fit best to the observed nfSFS at the mitochondrial locus. We omit the class of Dirac coalescents and algebraic growth models from further analysis since the observed frequency spectrum clearly does not fit to this model class. For each other model class used, we record the prior parameters from the 0.5% of the *n*_r_ = 200,000 simulations that have the smallest *ℓ*^2^ distance to the observed nfSFS (summary statistics). This gives an approximate sample of the posterior distribution of *π*(*α*| observed 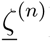) resp. *π*(*β*| observed 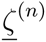). Again, we used the lumped nSFS as summary statistics. Figure S7 shows the posterior distributions for different lumping numbers.

**Figure S7:**
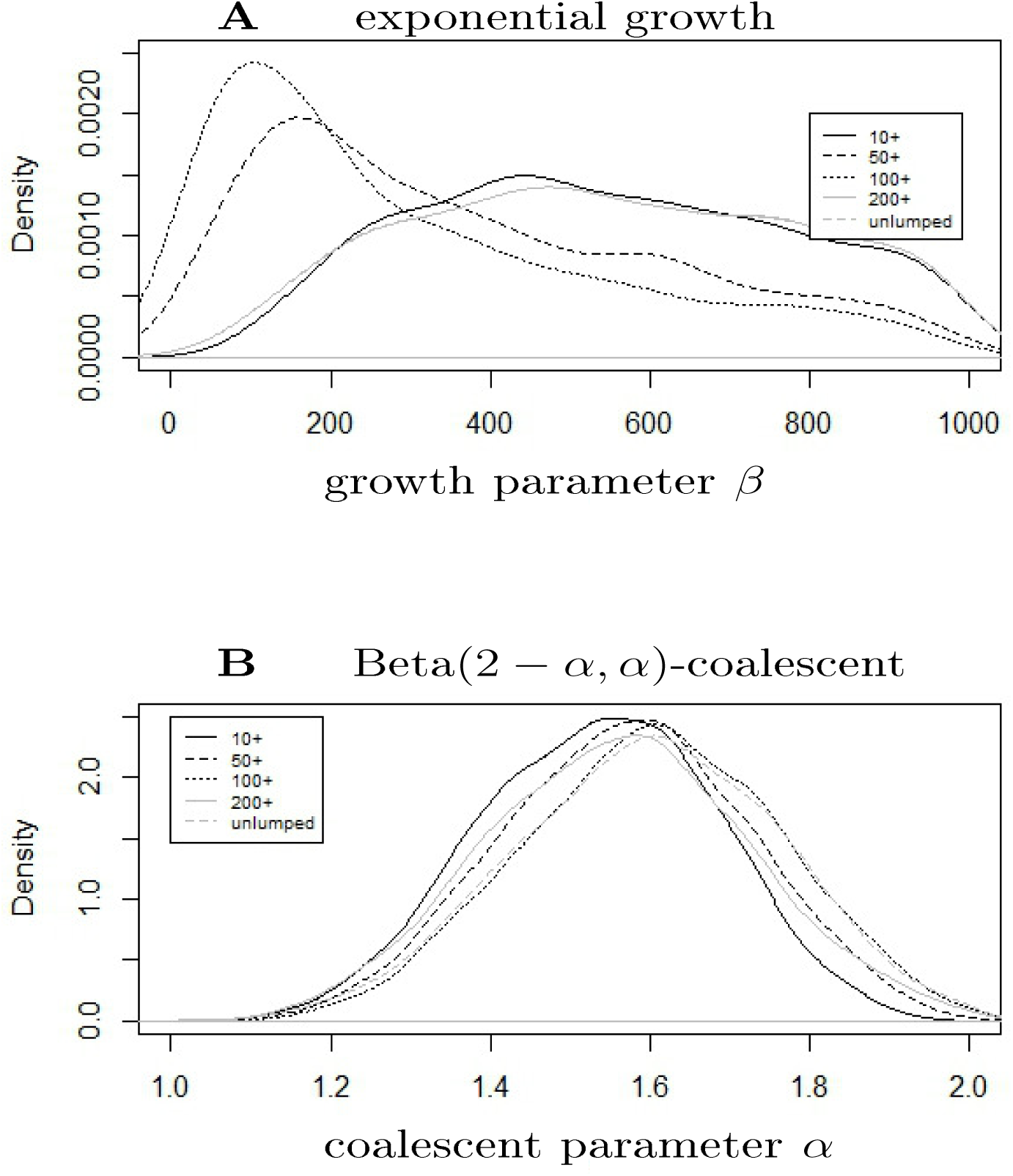
Approximate posterior density of the coalescent parameter from ABC fitting of the (**A**) growth, and (**B**) Beta *n*-coalescent model classes to the observed nfSFS in the Atlantic cod data. Denote by *α* the Beta *n*-coalescent parameter, *β* the growth rate. Priors were uniform on both sets.

### ABC quality control for the Árnason (2004) data

We follow the recommendation from the `R` package abc (Csilléry *et al,* 2012) and perform three checks of quality for the presented ABC approach. We focus on the lumping which gives the clearest distinction, namely the lumping of all classes with mutation counts 10 or higher (class 10+). All checks are performed using the `R` package abc

To assess the general ability to distinguish between the two model classes in the setting (i.e., number of observed mutations and sample size) given by the Atlantic cod mtDNA data from Árnason (2004), we again employ a leave-one-out cross-validation as described in Methods. See Tables S10 for the results.

**Table S10:** Approximations of the mean posterior probabilities and misclassification probabilities (based on *n*_cv_ = 12,000 cross-validations) for the ABC model comparison between models `E`, `A`, `B`, `D` for tolerance *x* = 0.005, sample size *n* = 1278 and mutation rate estimated via Watterson’s estimator from *s* = 39 observed mutations. The lumped nfSFS (10+) was used as summary statistics. The entries are listed as 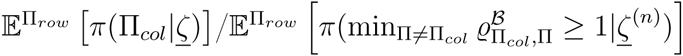

|  | E | A | B | D |
| --- | --- | --- | --- | --- |
| E | 0.79/0.88 | 0.00/0.00 | 0.21/0.12 | 0.00/0.00 |
| A | 0.00/0.00 | 0.24/0.25 | 0.06/0.01 | 0.70/0.74 |
| B | 0.24/0.18 | 0.01/0.00 | 0.71/0.79 | 0.03/0.02 |
| D | 0.00/0.00 | 0.03/0.03 | 0.08/0.03 | 0.90/0.94 |

To assess the quality to distinguish the parameters within one model class, we again use leave-one-out cross-validations (*n*_cv_ = 12,000). The parameter of each simulation chosen for cross-validation is estimated as the median of the 0.5% of simulations with the smallest *ℓ*^2^ distance to the chosen simulation. Figure S8 shows the resulting scatter plots of the parameters of the chosen simulations and the corresponding estimations.

**Figure S8:**
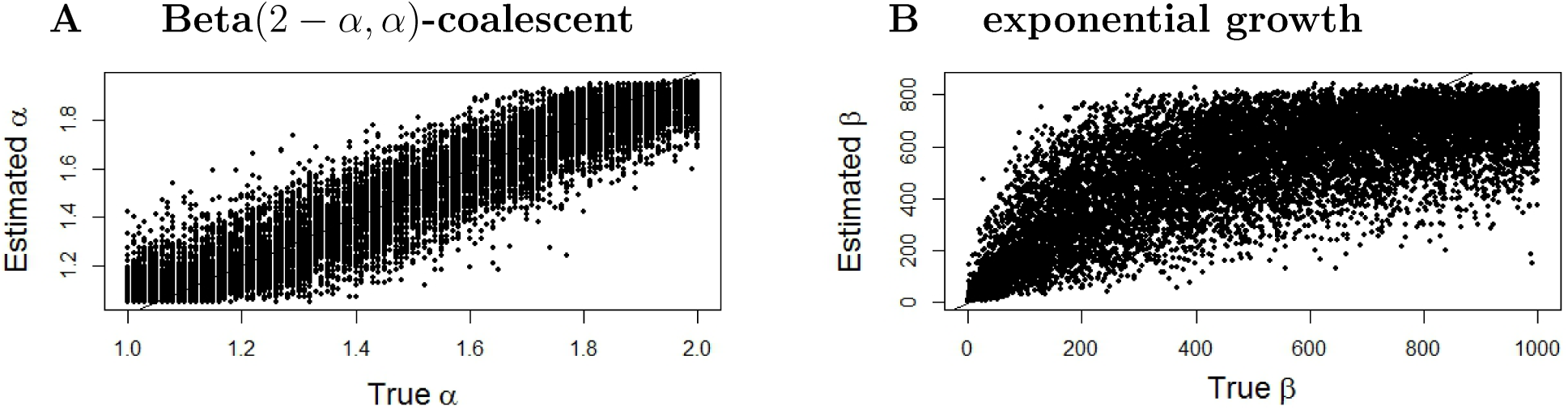
Scatter plots of estimated vs. true parameters of *n*_cv_ = 12000 cross-validated simulations in the (**A**) Beta coalescent; (**B**) exponential growth.

To see whether the posterior distributions given the cod mtDNA data from ÁRNASON (2004) define models under which the observed data is reproducible, we perfomed posterior predictive checks by simulating the 10+ lumped nfSFS under the posterior distribution (i.e., simulating once from each parameter set of each of the 1,000 accepted simulations) for each model class and compare these with the nfSFS observed. See Figure S9 for the results within each nfSFS class. To assess the minimal *l*^2^ distance of the simulations using the posterior parameter distributions from the observed nfSFS, we simulated 5 replications under the posteriors. The minimal *l*^2^ distance was 0.04 under the posterior growth model and 0.06 under the posterior Beta coalescent model.

**Figure S9:**
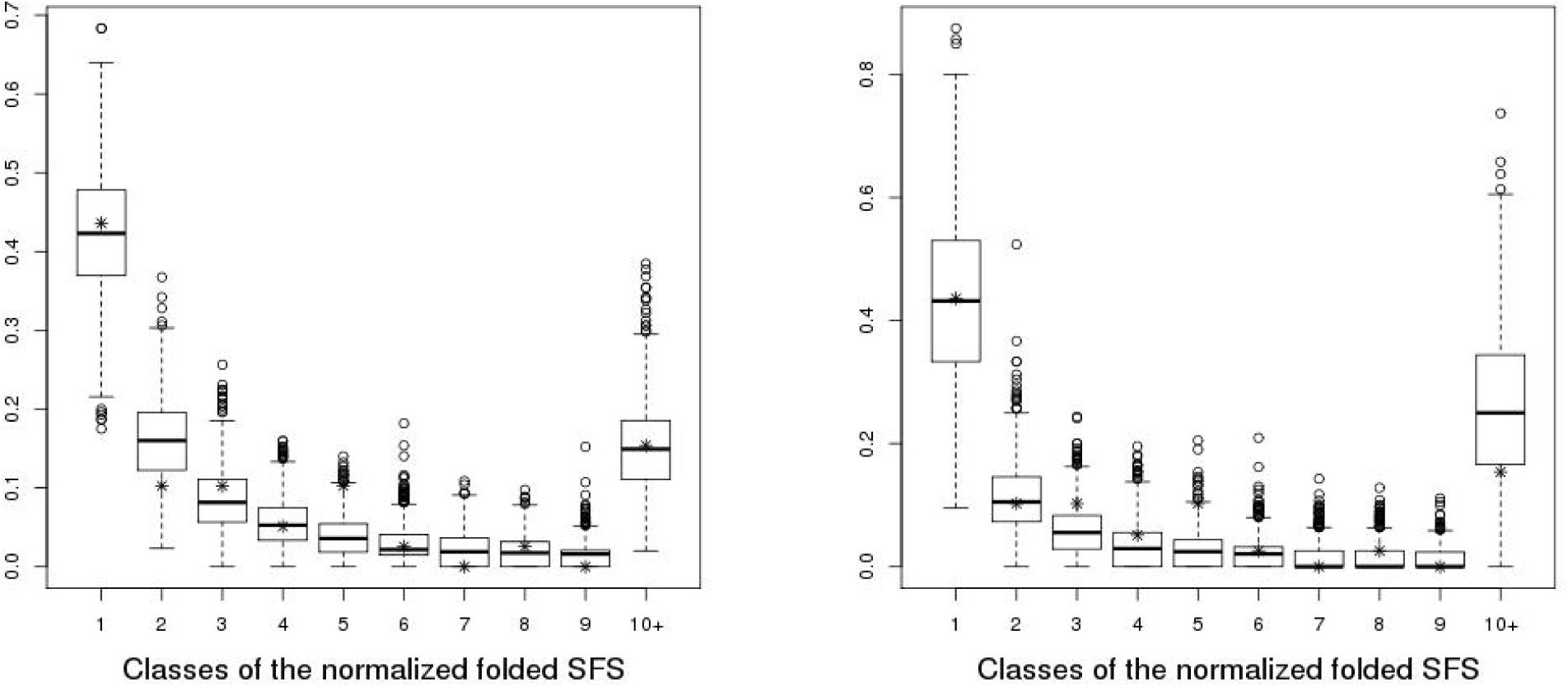
Posterior predictive checks with 1,000 simulations of the nfSFS under the approximate posterior distributions given the cod data from Árnason (2004) for the (**A**) Beta coalescent model class; (**B**) growth model class. Asterisks denote the observed values in the data.

The quality checks reveal that we can not distinguish well within the model classes of exponential growth and of Beta coalescents, but moderately between them. Additionally, the ABC approach distinguishes well between these two classes on one hand and the (nonfitting) other two classes `A`, `D`. The posterior predictive checks reveal that both model classes can produce the observed values in each class of the nfSFS, but do not match well in *l*^2^ to the actual observed nfSFS. Neither model class thus captures the observed nfSFS well.

## Footnotes

1 The beta-coalescent is also well-defined for *α* ∈ (0, 2], but we restrict to a smaller parameter range corresponding to the population model in Schweinsberg (2003).

